# Knockdown of Atg1 or Atg18 in adult adipocytes impairs lipophagy and reduces lifespan of fruit flies (*Drosophila melanogaster*)

**DOI:** 10.1101/2023.03.22.533865

**Authors:** Mariah Bierlein, Joseph Charles, Trevor Polisuk-Balfour, Heidi Bretscher, Micaela Rice, Jacklyn Zvonar, Drake Pohl, Lindsey Winslow, Brennah Wasie, Sara Deurloo, Jordan Van Wert, Britney Williams, Gabrielle Ankney, Zachary Harmon, Erica Dann, Anna Azuz, Alex Guzman-Vargas, Elizabeth Kuhns, Thomas P. Neufeld, Michael B. O’Connor, Felix Amissah, Changqi C. Zhu

## Abstract

Autophagy, a lysosome-based eukaryotic cellular degradation system, has previously been implicated in lifespan regulation in different animal models. In this report, we show that expression of the RNAi transgenes targeting the transcripts of the key autophagy genes such as *Atg1* or *Atg18* in adult fly muscle or glia does not affect the overall levels of autophagosomes in those tissues and does not change the lifespan of the tested flies, but lifespan reduction phenotype has become apparent when *Atg1* RNAi or *Atg18* RNAi is expressed in a non-tissue-specific manner through a *Tub-Gal4* in adult flies or after lipophagy is eradicated through the knockdown of Atg1 or Atg18 in adult fly adipocytes. Lifespan reduction was also observed when Atg1 or Atg18 was knocked down in adult fly enteroblasts and middle gut stem cells. Over-expression of wildtype Atg1 in adult fly muscle or adipocytes reduces lifespan. High levels of ubiquitinated protein aggregates could be the culprit of the reduced lifespan of Atg1 over-expression flies. Our research data presented here have highlighted the important functions of the key autophagy genes in adult fly adipocytes, enteroblasts, and midgut stem cells for lifespan regulation and their undetermined functions in adult fly muscle and glia.

**Summary:** In this research, we have demonstrated that the key autophagy genes play important roles in fly lifespan regulation through adult adipose tissues, enteroblasts, and middle gut stem cells.

## Introduction

Evolutionarily conserved autophagy in eukaryotic species has been implicated in the regulation of cellular energy homeostasis when cells are under nutrient-deprived conditions as well as in the clearance of defective cytoplasmic organelles such as damaged mitochondria and endoplasmic reticulum (Mizushima, 2007; Takeshige et al., 1992). In addition, autophagy has been found to play important roles in protein secretion, intracellular vesicle trafficking, endocytosis, and lipophagy in eukaryotes (Schott et al., 2022; Shravage et al., 2013; Zhang et al., 2019). These diverse and yet related cellular functions are regulated through the proteins encoded by more than twenty autophagy-related genes (Atg). Among the well-conserved *Atg* genes from different eukaryotic species, *Atg1*, which encodes a serine and threonine kinase, appears to be the most potent player in helping initiate the autophagy process, through which double-layered phospholipid membranes are recruited via membrane-associated receptors to form autophagosomes, which then deliver cellular contents to the lysosomes for degradation (Mizushima, 2007).

Autophagy is known to be regulated by the cellular growth and protein synthesis regulators of target of rapamycin (TOR) pathway from fungal yeast to fruit flies and to mammals (Noda and Ohsumi, 1998; Scott et al., 2004; Hosokawa et al., 2009). In fungal yeast, TOR inhibitor rapamycin induces autophagy independent of starvation (Noda and Ohsumi, 1998). Similar regulatory mechanism of autophagy by TOR signaling exists in fruit flies. Defective TOR signaling induces autophagy even when the animals are under well-fed conditions (Scott et al., 2004). Conversely, autophagy can be effectively suppressed in the starved cells when TOR signaling is artificially maintained by using genetic means either through the forced expression of a wild type PI3 kinase (PI3K) or through the expression of a wild type Rheb gene, which encodes a GTP-binding protein and works as a positive regulator of TOR signaling (Scott et al., 2004). The regulation of autophagy by TOR signaling is achieved through protein-protein interactions of one of its two TOR protein complexes, mTORC1 protein complex, which interacts with the ULK1 (mammalian Atg1 homologue protein)-Atg13-FIP200 protein complex. mTORC1 phosphorylates ULK1 and Atg13 in a nutrient-dependent manner in mammalian cells (Hosokawa et al., 2009).

Being a serine/threonine kinase, Atg1 exhibits multiple functions in eukaryotic cells, which include autophagy induction and cell growth inhibition, both of which seem to be interrelated to make cells conserve as much energy as they can to stay alive under starved conditions. Cell growth inhibition by Atg1 has been demonstrated to be achieved through its regulation of Tor/S6 kinase pathway and/or Yorkie function (Lee et al., 2007; Tyra et al., 2020). For normal cellular growth and energy homeostasis, the antagonizing function of TOR signaling and autophagy seems to be an important molecular mechanism employed by eukaryotic cells. Other autophagy related genes include *Atg5*, *Atg7, Atg8*, *Atg9*, *Atg12, Atg14, Atg16, Atg17*, and *Atg18*. The proteins encoded by *Atg5, Atg12,* and *Atg16* form an Atg12-Atg5-Atg16 protein complex, which associates with Atg1 protein via Atg12. The whole protein complex serves as a scaffold for the membrane-based phagophore to form (Harada et al., 2019). Atg1 kinase acts early in autophagosome formation by phosphorylating Atg9 protein (Papinski et al., 2014). Phosphorylated transmembrane protein Atg9 is required for the efficient recruitment of Atg8 and Atg18 proteins to the site of forming autophagosomes (Papinski et al., 2014). Atg9 (presumably the phosphorylated form) is known to be involved in conducting lipid transport from cytoplasmic membrane pools to the pre-autophagosomal structure, which will eventually lead to the formation of mature autophagosomes (Mari et al., 2010; Rao et al., 2016; Suzuki et al., 2015; Tooze, 2010; Yamamoto et al., 2012). Both Atg9 and lipid-binding protein Atg18 have been demonstrated to be required for autophagy activities in fungal yeast (He et al., 2008; Obara et al, 2008). The role of Atg18 in autophagosome formation appears to be achieved through its common membrane tubulation and scission functions (Gopaldass et al., 2017). The degradation of the cellular components enclosed in the autophagosomes takes place in lysosomes after fusion of these two membrane-bound organelles.

At the tissue and organismal level, autophagy has been identified to affect adult tissue homeostasis and normal lifespan. In mammals, Beclin 1, one of the key autophagy proteins, has been discovered as a potent tumor suppressor (Qu et al., 2003; Yue et al., 2003). In fruit flies, germline mutations of individual Atg7 and Atg9 genes affect autophagy and adult lifespan (Juhász et al., 2007; Wen et al., 2017). Similar lifespan defect was observed in Atg7 conditional knockout mice (Komatsu et al., 2006). Over-expression of some key autophagy genes such as Atg5 and Atg8a appeared to extend lifespan in mice and fruit flies, respectively (Pyo et al., 2013; Simonsen et al., 2008). For these lifespan results obtained either through loss-of-function studies of the mutant animals derived from germline mutations or through over-expression of individual wild type Atg transgenes in both development and adult life, a question about whether autophagy-mediated lifespan regulation is achieved through developmental regulation or through the regulation of specific adult tissues or a combination of both by autophagy genes has not previously been addressed.

To better understand the tissue specific requirement of the key autophagy genes in adult fruit fly, in our present study we used a Gal80 temperature sensitive protein (Gal80^ts^)-based Gal4 TARGET gene expression system (McGuire et al., 2004) to assess how several autophagy genes affect adult fly lifespan by selectively suppressing autophagy in specific adult tissues. We also over-expressed either a wild type *Atg1* transgene or a wild type *Atg8a* transgene in adult fruit fly muscle and adipose tissue to examine the potential lifespan changing effects by the two genes. Our experimental data showed that expression of *Atg1 RNAi* or *Atg18 RNAi* in adult fly muscle or glial cells does not alter the overall levels of the autophagosomes, nor does it change the lifespan when compared to the controls. However, the requirement of the key autophagy genes in adult fly lifespan regulation was demonstrated when a non-tissue-specific Gal4 driver *Tub-Gal4* fly line, an adipose tissue and glial cell Gal4 driver *Daw-Gal4* fly line, and a midgut enteroblast and stem cell Gal4 driver *Esg-Gal4* fly line were used to induce the knockdown effects for *Atg1* or *Atg18* genes in adult flies. In contrast to some of the previously published findings, our experimental results have demonstrated that over-expression of wild type *Atg1* but not *Atg8a* in adult fly muscle or adipose tissue reduces lifespan. Under no circumstances did we observe any lifespan extension effect by over-expressing wild-type *Atg1* or *Atg8a* in adult fly muscle or adipose tissue. The lifespan reduction effect elicited by over-expression of *Atg1* in adult fly muscle correlates with the high levels of protein aggregates induced by this transgene in those tissues.

## Results

### Characterization of autophagy in adult fruit flies through a UAS-mCherry-Atg8a fly line

To assess the potential roles of autophagy in adult fly lifespan regulation, we first monitored the levels of autophagosomes in adult muscles, adipocytes, and glia of control, *Atg1 RNAi*, and *Atg18 RNAi* expression flies with a *UAS-mCherry-Atg8a* fly line (Chang and Neufeld, 2009) and a muscle-specific Gal4 fly line, *Mhc-Gal4*, and a glia and adipocyte Gal4 fly line, *Daw-Gal4*. For this and the rest of the study, we adopted the *Tub-Gal80^ts^*-based Gal4 TARGET gene-switch system (McGuire et al., 2004) to induce adult tissue-specific expression of any transgenes that are under the control of yeast upstream activation DNA sequences (UAS) with a temperature shift for the culture of newly hatched flies from 18°C to 29°C. All the fruit fly crosses involving *Tub-Gal80^ts^* transgene and other specific Gal4/UAS transgenes were carried out at 18°C so that Gal80^ts^ can actively suppress Gal4 function, and the normal development of the flies can proceed. After hatching from 18°C, the flies inherited a copy of *Tub-Gal80^ts^* transgene and other specific Gal4/UAS transgenes were switched to a permissive temperature (29°C) as adults, in which Gal80^ts^ is inactive, thus allowing Gal4-induced expression of any specific transgenes of UAS control. By using the described experimental strategy, fruit flies harboring different transgenes as illustrated in Fig. 1A-I were examined for the presence or absence of mCherry-Atg8a- labeled autophagosomes (red puncta). Different levels of autophagosomes as marked by the mCherry-Atg8a fusion protein were observed in adult fly indirect flight muscles (Laurichesse and Soler, 2020), glial cells, and adipocytes of control flies (Fig. 1A, 1D, & 1G) as well as in the muscles and glial cells of *Atg1 RNAi* or *Atg18 RNAi*-expressing flies (Fig. 1B, 1C, 1E, & 1F). There is no obvious difference of mCherry-Atg8a-labeled autophagosomes in the muscles or glial cells of control and *Atg1 RNAi* or *Atg18 RNAi*-expressing flies (Fig. 1B vs 1A, 1C vs 1A, 1E vs 1D, and 1F vs 1D). However, co-expression of either *UAS-Atg1 RNAi* or *UAS-Atg18 RNAi* with *UAS-mCherry-Atg8a* via *Daw-Gal4* in adipocytes clearly eliminated all mCherry-Atg8a-labeled autophagosomes as shown in Fig. 1H and 1I, respectively. The lack of autophagy phenotype in adult fly muscle and glia could be due to the ineffective knockdown of the transcripts of *Atg1* and *Atg18* by the expression of *Atg1 RNAi* and *Atg18 RNAi* in those tissues as shown in Fig. 1J (p>0.05). Alternatively, Atg1-independent autophagy may play a role in adult fly muscle and glial cells, which we can’t rule out now.

**Figure 1.**
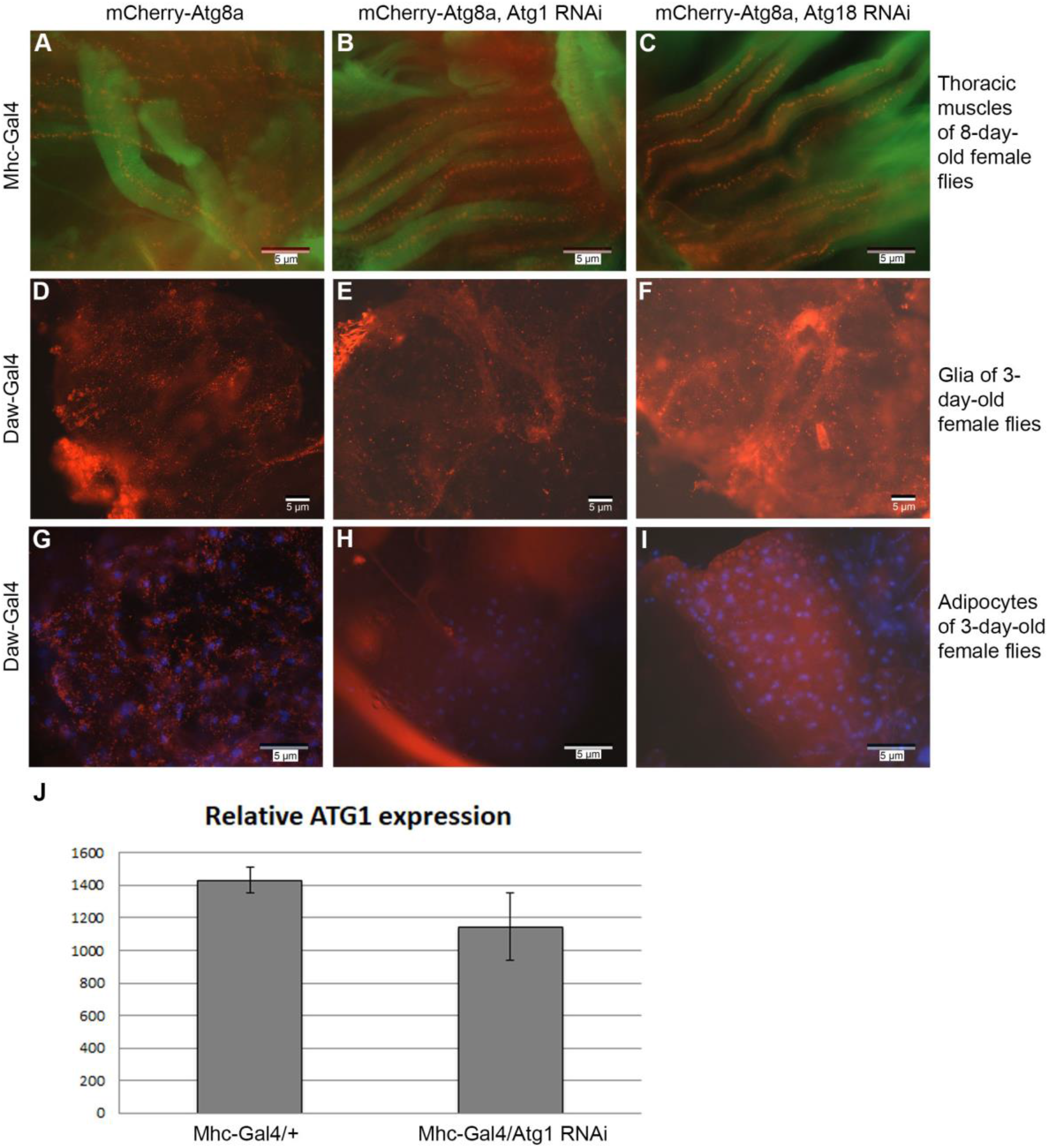
mCherry-Atg8a-labeled autophagosomes in adult fly muscle and glia of control and *Atg1 RNAi*- and *Atg18 RNAi*-expressing fruit flies and their absence from the adipocytes of Atg1 or Atg18 adipocyte-specific knockdown flies. **(A-C)** mCherry-Atg8a-labeled autophagosomes (red puncta) can be clearly identified in the live indirect flight muscles of an 8-day-old female control fly with a genotype of “*Tub-Gal80^ts^/UAS-mCherry-Atg8a, Mhc-Gal4/+*” (A) and in the live muscles of an 8-day-old *Atg1 RNAi*-expressing female fly of a genotype “*Tub-Gal80^ts^/UAS-mCherry-Atg8a, Mhc-Gal4/UAS-Atg1 RNAi*” (B) and in the live muscles of an 8-day-old *Atg18 RNAi-expressing* female fly with a genotype “*Tub-Gal80^ts^/UAS- mCherry-Atg8a, Mhc-Gal4/UAS-Atg18 RNAi*” (C). Some red puncta of picture (A) that were not part of the muscle tissues could have been caused by the separation of autophagosomes from the thoracic indirect flight muscles because of imprecise dissection of the tissues with forceps. **(D-F)** No obvious differences are seen for the levels of the mCherry-Atg8a-labeled autophagosomes from the glial cells of the central brains of a 3-day-old control female fly with a genotype of “*Tub-Gal80^ts^/UAS- mCherry-Atg8a, Daw-Gal4/+*” (D), a 3-day-old glial *Atg1 RNAi-*expressing fly with a genotype of “*Tub-Gal80^ts^/UAS-mCherry-Atg8a, Daw-Gal4/UAS-Atg1i*” (E), and a 3-day-old glial *Atg18 RNAi-*expressing fly with a genotype of “*Tub-Gal80^ts^/UAS-mCherry-Atg8a, Daw-Gal4/UAS-Atg18i*” (F). **(G-I)** Knockdown of Atg1 (H) or Atg18 (I) in adipocytes clearly eliminated mCherry-Atg8a-labeled autophagosomes when compared to the control in (G). (G) Adipocytes of a 3-day-old control female fly with a genotype of “*Tub-Gal80^ts^/UAS-mCherry-Atg8a, Daw-Gal4/+*” are full of mCherry-Atg8a-labeled autophagosomes, but mCherry-Atg8a-labeled autophagosomes can’t be identified in the adipocytes of a 3-day-old *Daw-Gal4*-induced Atg1 knockdown female fly with a genotype of “*Tub-Gal80^ts^/UAS-mCherry-Atg8a, Daw-Gal4/UAS-Atg1 RNAi*” (H) and a 3-day-old *Daw-Gal4*-induced Atg18 knockdown female fly with a genotype of “*Tub-Gal80^ts^/UAS-mCherry-Atg8a, Daw-Gal4/UAS-Atg18 RNAi*” (I). All the flies used in the experiments described here were raised at 29°C since their hatching from 18°C. The live muscles from (A) to (C) were labeled by CytoPainter Phalloidin-iFlor 488 (from Abcam) for F-actin in green. The live adipose tissues from (G) to (I) were labeled by 4 ′, 6-diamidino-2-phenylindole (DAPI) for nuclei in blue. **(J)** Comparison for the relative expression levels of the *Atg1* mRNA molecules isolated from the control flies with a genotype of “*Tub-Gal80^ts^/UAS-mCherry-Atg8a, Mhc-Gal4/+”* simplified as “Mhc-Gal4/+” and *Mhc-Gal4*-induced *Atg1 RNAi*-expressing flies with a genotype of “*Tub-Gal80^ts^/UAS-mCherry-Atg8a, Mhc-Gal4/UAS-Atg1 RNAi*” labeled as “Mhc-Gal4/Atg1 RNAi” through qPCR. The flies of the same age used in (J) were raised at 29°C when the total RNA was extracted for qPCR.

### Expression of Atg1 RNAi, Atg5 RNAi, Atg9 RNAi, or Atg18 RNAi through a Tub-Gal4 fly line in adult flies shortens lifespan

*Atg1*, *Atg5*, *Atg9*, and *Atg18* are four key autophagy genes whose functions in autophagy induction in fruit flies had previously been demonstrated by other researchers (Nagy et al., 2014; Scott et al., 2004; Tang et al., 2013). By mating individual RNAi transgenic fly lines including a *UAS-Atg1 RNAi* (*Atg1i*) line, a *UAS- Atg5 RNAi* (*Atg5i*) line, a *UAS-Atg9 RNAi* (*Atg9i*) line, or a *UAS-Atg18 RNAi* (*Atg18i*) line to a compound transgenic fly line “*Tub-Gal80^ts^, Tub-Gal4/Tm6^TbHu^*” at 18°C and collecting newly hatched male and female flies that inherited a copy of *UAS-Atg1i, UAS-Atg5i, UAS-Atg9i,* or *UAS-Atg18i* and a copy of *Tub-Gal80^ts^* and *Tub-Gal4* in separate vials and raising them at 29°C together with the control flies, we discovered that *Tub-Gal4*-induced knockdown of the transcripts of *Atg1, Atg5, Atg9,* or *Atg18* shortened the mean and maximum lifespans of both male and female flies when compared to the controls (Fig. 2A-F, and Supplemental table 1 to 4). *Tub-Gal4* can drive the expression of any transgene that’s under the control of *UAS* in many different fly tissues, and therefore we can’t conclude from these experiments which adult fly tissue is required for lifespan regulation through autophagy genes, but with these few experiments we are certain that autophagy genes are required in adult flies for their lifespan regulation. For the *Tub-Gal4*-driven Atg1 knockdown female flies, the 29.9-day mean lifespan was about 13.5 and 17.5 days shorter than the mean lifespans of the two control groups of flies, respectively (Fig. 1A, p<0.05, and Supplemental table 1). The 45-day maximum lifespan of those *Tub-Gal4*-driven Atg1 knockdown female flies was 10 and 12 days shorter than the maximum lifespans of the two control groups of flies, respectively (Fig. 1A, and Supplemental table 1). The mean and maximum lifespans of *Tub-Gal4*-driven Atg1 knockdown male flies were significantly reduced too when compared to the controls (Fig. 1B, and Supplemental table 2). The combined lifespan data from the *Tub-Gal4*-driven Atg1 knockdown adult female and male flies (Fig. 2C) just showed the same reduction trend as those individual sets of data from *Tub-Gal4*-driven Atg1 knockdown adult female or male flies when analyzed separately. Similar lifespan reduction phenotypes were observed for the *Tub-Gal4*-driven knockdown of Atg5, Atg9, or Atg18 in adult flies (Fig. 2D-2F, and Supplemental table 3 and 4). Because of the concerted functions of these key Atg genes in autophagy induction, it’s not surprising to see consistent lifespan reduction phenotypes when the transcripts of these genes were knocked down separately and in a non-tissue-specific manner in adult flies.

**Figure 2.**
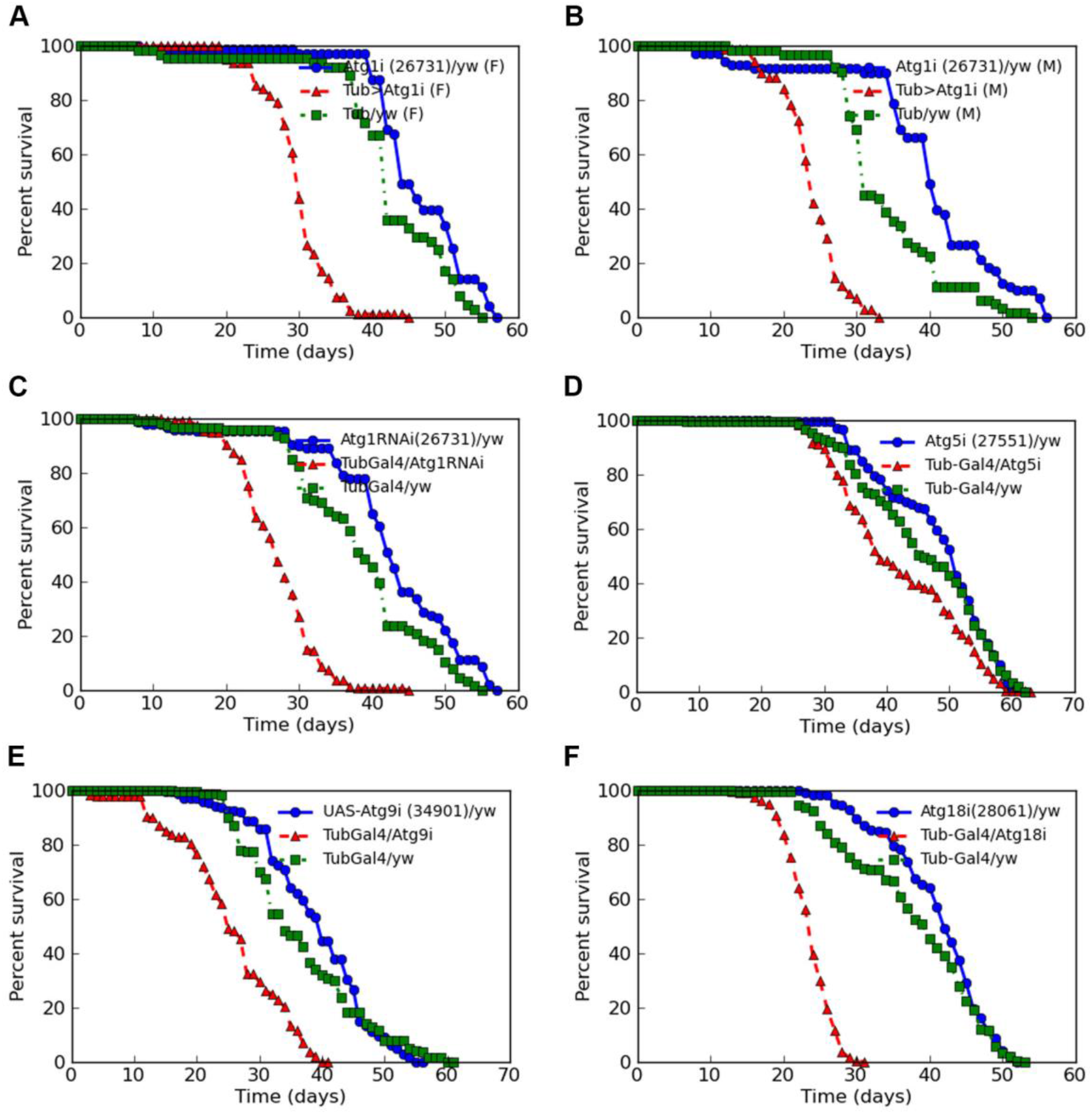
*Tub-Gal4*-induced knockdown of Atg1, Atg5, Atg9, or Atg18 in adult flies reduced lifespan. **(A)** The lifespan shortening effect caused by the expression of a *UAS-Atg1 RNAi* transgene (Bloomington stock 26731) through a non-tissue-specific Gal4 driver *Tub-Gal4* in adult female flies at 29°C. The Atg1 knockdown female flies with a genotype of “*Tub-Gal80^ts^, Tub-Gal4/UAS-Atg1 RNAi*” (labeled as Tub>Atg1i (F)) cultured at 29°C had a mean lifespan of 29.9 days and a maximum lifespan of 45 days (red curve), whereas the two control groups of female flies with a genotype of “*UAS-Atg1 RNAi/yw*” (labeled as Atg1i(26731)/yw (F), blue curve) or “*Tub-Gal80^ts^, Tub-Gal4/yw*” (labeled as Tub/yw (F), green curve) from 29°C showed a 46.31-day and a 42.33-day mean lifespan, a 57-day and a 55-day maximum lifespan, respectively (*p<0.05*). **(B)** *Tub-Gal4-*induced knockdown of Atg1 at 29°C also significantly shortened the mean and maximum lifespans of male flies with a genotype of “*Tub-Gal80^ts^, Tub-Gal4/UAS-Atg1 RNAi*” (labeled as Tub>Atg1i (M), red curve) when compared to their controls (labeled as Atg1i(26731)/yw (M) and Tub/yw (M), blue and green curves) (*p<0.05*). **(C)** Comparison of the combined survivorships of *Tub-Gal4-*induced Atg1 knockdown female and male flies to their controls (red versus green and blue curves). The reduced survivorship from the combined *Tub-Gal4-*induced Atg1 knockdown female and male flies is obvious when compared to the combined survivorships of the control female and male flies (*p<0.05*). **(D)** Knockdown of Atg5 through *Tub-Gal4* also reduced the overall lifespan of the tested male and female flies when compared to the controls. **(E-F)** The lifespan reduction effect was obvious when either Atg9 (E) or Atg18 (F) was knocked down through *Tub-Gal4* at 29°C in adult flies (red survivorship curves for Atg9 knockdown flies or Atg18 knockdown flies versus the blue and green survivorship curves of control flies).

### Expression of Atg1 RNAi, Atg5 RNAi, Atg9 RNAi, or Atg18 RNAi in adult fly muscle does not alter the overall lifespans of the tested flies

Although the expression of *Atg1 RNAi* or *Atg18 RNAi* in adult fly muscle did not alter the overall levels of mCherry-Atg8a-labeled autophagosomes (Fig. 1A to 1C), the fact that *Tub-Gal4*-induced expression of either transgene can shorten the lifespan of both male and female flies significantly (Fig. 2A to 2C and Fig. 2F) prompted us to test if the expression of these same transgenes in adult fly muscle can change the lifespan or not. For this purpose, we mated individual *UAS-Atg1 RNAi* fly line, *UAS-Atg5 RNAi* line, *UAS-Atg9 RNAi* line, or *UAS-Atg18 RNAi* line to a “*Tub-Gal80^ts^, Mhc-Gal4/Tm6^TbHu^*” fly line at 18°C and cultured their progenies that inherited a copy of each RNAi transgene and a copy of “*Tub-Gal80^ts^, Mhc-Gal4”* together with control flies at 29°C for lifespan observation. The control flies were generated by mating each of the parental fly lines for the experimental flies to a yellow white fly line (*yw*) under the same conditions as used for the experimental groups of flies. We found that consistent with the autophagy results observed in Fig. 1A-C, the expression of none of the *Atg1 RNAi*, *Atg5 RNAi, Atg9 RNAi,* or *Atg18 RNAi* transgenes in adult female and male fly muscles can change the overall lifespans of the tested flies when compared to the controls (Fig. 3A-F, Supplemental table 5 and 6). Starvation-induced autophagy provides cells with needed nutrients and energy (Noda and Ohsumi, 1998; Scott et al., 2002). To test if the expression of *Atg1 RNAi* in adult fly muscles can impair the normal survivorship of the adult flies under starved conditions, we starved *Atg1 RNAi*-expressing flies and their controls and showed that the expression *Atg1 RNAi* in adult fly muscle under starved conditions does not enhance the lethality rate of these flies when compared to the controls (Fig. 3G-H).

**Figure 3.**
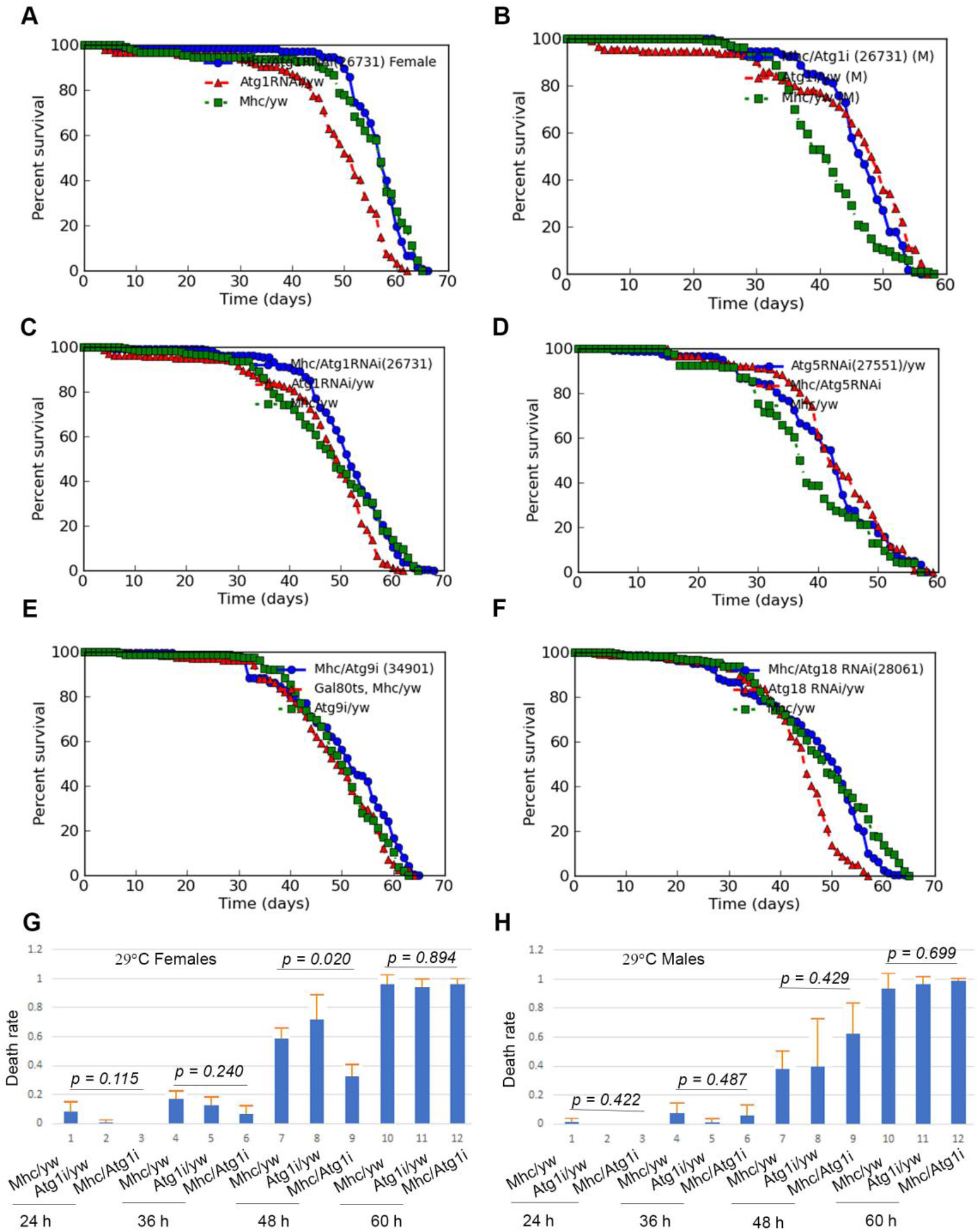
The expression of *Atg1 RNAi*, *Atg5 RNAi*, *Atg9 RNAi*, or *Atg18 RNAi* in adult fly muscle at 29°C does not change the overall lifespan of these flies when compared to the controls. **(A)** The survivorships of the adult muscle-specific *Atg1 RNAi*-expressing female flies (*Mhc-Gal4/UAS-Atg1 RNAi (26731)* and their controls (Atg1RNAi/yw and Mhc/yw). No statistically significant difference is seen for the mean and the maximum lifespans of the muscle-specific *Atg1 RNAi- expressing* female flies and their controls (blue curve versus green curve, *p>0.05*). **(B)** Adult muscle-specific *Atg1 RNAi*-expressing male flies did not show any obvious lifespan reduction phenotype either when compared to the controls (blue curve versus red curve, *p>0.05*). **(C)** Comparison of the combined lifespans of adult muscle-specific *Atg1 RNAi*-expressing female and male flies to their controls. No apparent lifespan shortening effect from *Atg1 RNAi* expression in adult fly muscle can be seen (Blue versus red and green curves). **(D-F)** The mean and maximum lifespans of the combined male and female flies with *Atg5 RNAi*, *Atg9 RNAi*, or *Atg18 RNAi* expression in adult fly muscles are not statistically different from those of the control flies (*p>0.05*). **(G-H)** Expression of *Atg1 RNAi* in adult male or female fly muscle (*Mhc/Atg1i*) does not negatively affect the overall starvation-afflicted survivorships of the tested flies when compared to the controls (*Mhc/yw* and *Atg1i/yw*). Instead, the female flies with *Atg1RNAi* expressed in adult fly muscle (*Mhc/Atg1i*) seemed to survive the starvation even better than the control flies (*Mhc/yw* and *Atg1i/yw*) 48 hours since the start of the starvation (G) (*p<0.02*). Orange bars represent standard deviations from ANOVA statistic test.

### Expression of Atg1 RNAi or Atg18 RNAi in adult fly adipose tissue or enteroblasts and gut stem cells but not in glial cells shortens lifespan

Lack of apparent lifespan phenotype when the RNAi transgenes against the key autophagy genes were expressed in adult fly muscle led us to test if the expression of *Atg1 RNAi* or *Atg18 RNAi* in adult fly glial or adipose tissue or enteroblasts and gut stem cells could alter the lifespans of the flies. For this purpose, we used a glial cell-specific Gal4 fly line, *Repo-Gal4*, and a glial and fat body Gal4 fly line, *Daw-Gal4,* to induce the expression of *Atg1 RNAi* or *Atg18 RNAi* in adult fly glial and adipose tissues, and an *Esg-Gal4* for enteroblasts and gut stem cells. As shown in Fig. 4A-B, expression of *Atg1 RNAi* or *Atg18 RNAi* in glial cells doesn’t change the overall lifespan when compared to the controls, which is consistent with the unchanged levels of mCherry-Atg8a-labeled autophagsomes in the glial cells from the flies that expressed either *Atg1 RNAi* or *Atg18 RNAi* through *Repo-Gal4* in those cells of adult brains (Fig. 1E-F vs 1D), but significant lifespan reduction can be observed when either *Atg1 RNAi* or *Atg18 RNAi* is expressed in enteroblasts and gut stem cells through *Esg-Ga4* (Fig. 4C- D) or in glia and fat body through *Daw-Gal4* (Fig. 4E-H).

**Figure 4.**
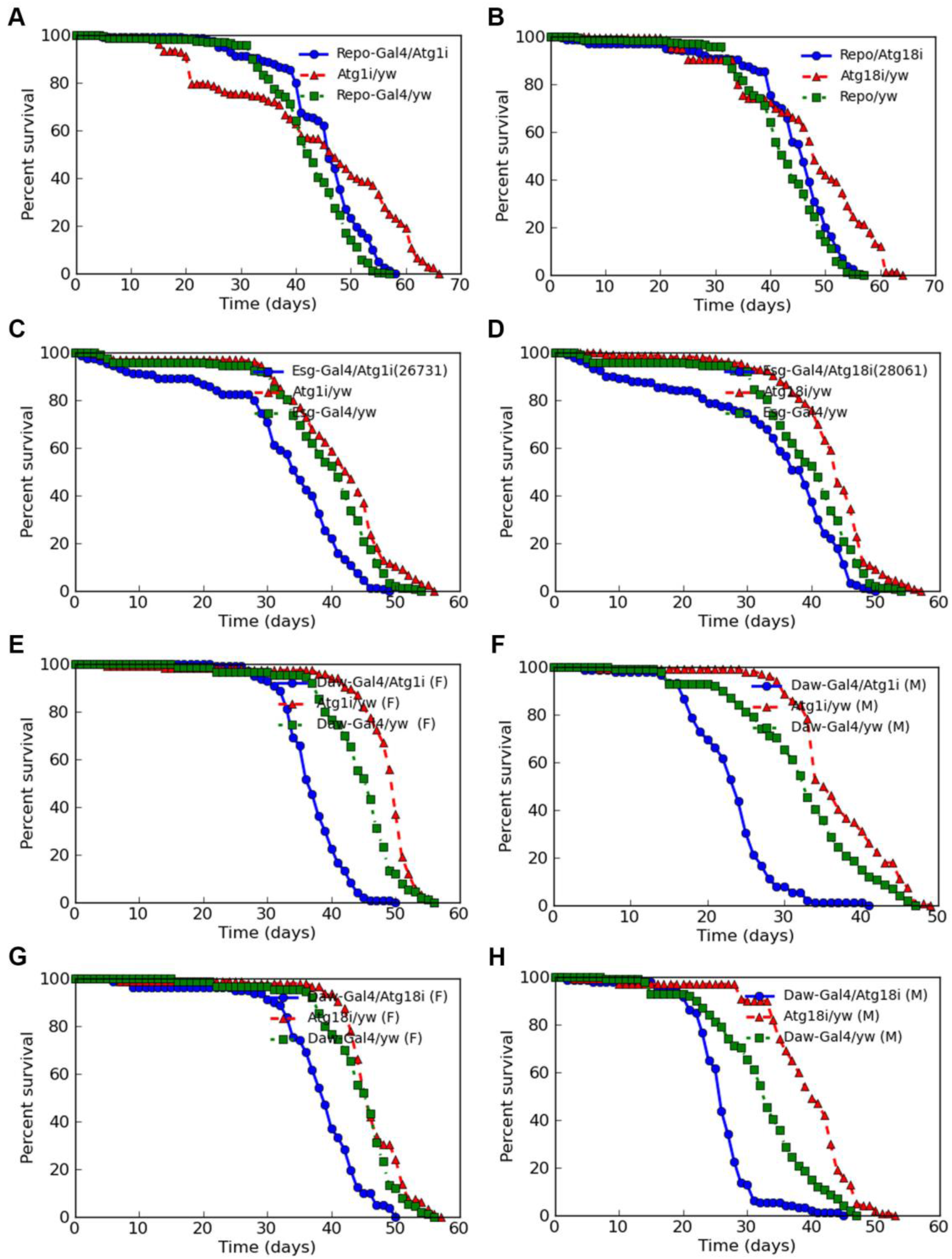
Expression of Atg1 RNAi or Atg18 RNAi in adult fly enteroblasts and gut stem cells or fat body tissues but not in glial cells shortens fly lifespan. **(A)** The expression of *Atg1* RNAi in adult fly glial cells (*Repo-Gal4/Atg1i*) does not alter the overall lifespans of the tested flies when compared to the controls (*Atg1i/yw* and *Repo-Gal4/yw*, *P>0.05*). **(B)** No lifespan alteration was observed when *Atg18* RNAi was expressed in adult fly glial cells either (Repo/Atg18i versus Atg18i/yw and Repo/yw, *P>0.05*). **(C-D)** Knockdown of Atg1 or Atg18 in adult male and female fly enteroblasts and gut stem cells at 29°C by using an *Esg-Gal4* fly line reduced the mean and the maximum lifespan of the tested flies when compared to the controls. **(C)** The combined mean lifespan of 32 days of the male and female flies with a genotype of “*Esg-Gal4/Tub-Gal80^ts^, UAS-Atg1 RNAi*” (*Esg-Gal4/Atg1i,* blue curve) for Atg1 knockdown in adult enteroblasts and gut stem cells at 29°C was about 6 and 8 days shorter than those of the control flies either having a genotype of “*Tub-Gal80^ts^, UAS-Atg1 RNAi/yw”* (*Atg1i/yw*, red curve) or a genotype of “*Esg-Gal4/yw”* (green curve), respectively (*p<0.00002*). The combined maximum lifespan of 49 days of the male and female flies with Atg1 knocked down in enteroblasts and adult gut stem cells was about 5 and 7 days shorter than those of the controls. **(D)** The combined 33.8 days mean lifespan of the male and female flies with Atg18 knocked down in adult fly enteroblasts and gut stem cells was about 5 and 9 days shorter than those of the control flies (blue versus red and green curves) (*p<0.05*). **(E-H)** Knockdown of Atg1 or Atg18 at 29°C by using a *Daw-Gal4* reduced both mean and maximum lifespans of the tested male and female flies when compared to the controls. **(E)** The 37-day mean lifespan of female flies raised at 29°C with Atg1 knocked down in adult adipose tissue of the flies with a genotype of “*UAS-GFP/Tub-Gal80^ts^, Daw-Gal4/UAS- Atg1 RNAi*” (*Daw-Gal4/Atg1i*, blue curve) was about 7 and 11 days shorter than those of the control flies with a genotype of “*UAS-GFP, Daw-Gal4/yw*” (*Daw-Gal4/yw*, green curve) and flies of “*Tub-Gal80^ts^, UAS-Atg1i/yw*” genotype (*Atg1i/yw*, red curves), respectively (*p<0.05*). **(F)** A similar lifespan reduction was observed for male flies with Atg1 knocked down in adult adipose tissue too. **(G)** Atg18 knockdown in adipose tissue of adult female flies through *Daw-Gal4* caused both mean and maximum lifespan reduction (blue curve versus green and red curves, *p<0.05*), which is very similar to the effect of Atg1 knockdown through *Daw-Gal4* in adult female flies as shown in (E). **(H)** The lifespan reduction effect caused by Atg18 knockdown in adipose tissue of adult male flies through *Daw-Gal4* is quite evident (blue curve versus green and red curves, *p<0.05*).

To know if the reduced lifespan of the flies with *Daw-Gal4-*induced expression of *Atg1 RNAi* or *Atg18 RNAi* is a precocious aging phenotype, we performed a typical aging phenotype assay by following the well-established negative geotaxis protocol for fruit flies (Gargano et al., 2005;). As demonstrated in Fig. 5A-L, *Daw-Gal4*-induced Atg1 or Atg18 knockdown female and male flies had demonstrated reduced vial wall climbing ability as they got old when compared to their corresponding control flies (Fig. 5D versus 5C, 5H versus 5G, and 5J versus 5I) (*p<0.05*). These vial wall climbing differences were not observed for 1-day-old *Daw-Gal4*-induced Atg1 or Atg18 knockdown and their control flies (Fig. 5A-B, 5E-F, and 5K-L), suggesting that the vial wall climbing phenotypes of Atg1 or Atg18 knockdown flies are age-related.

**Figure 5.**
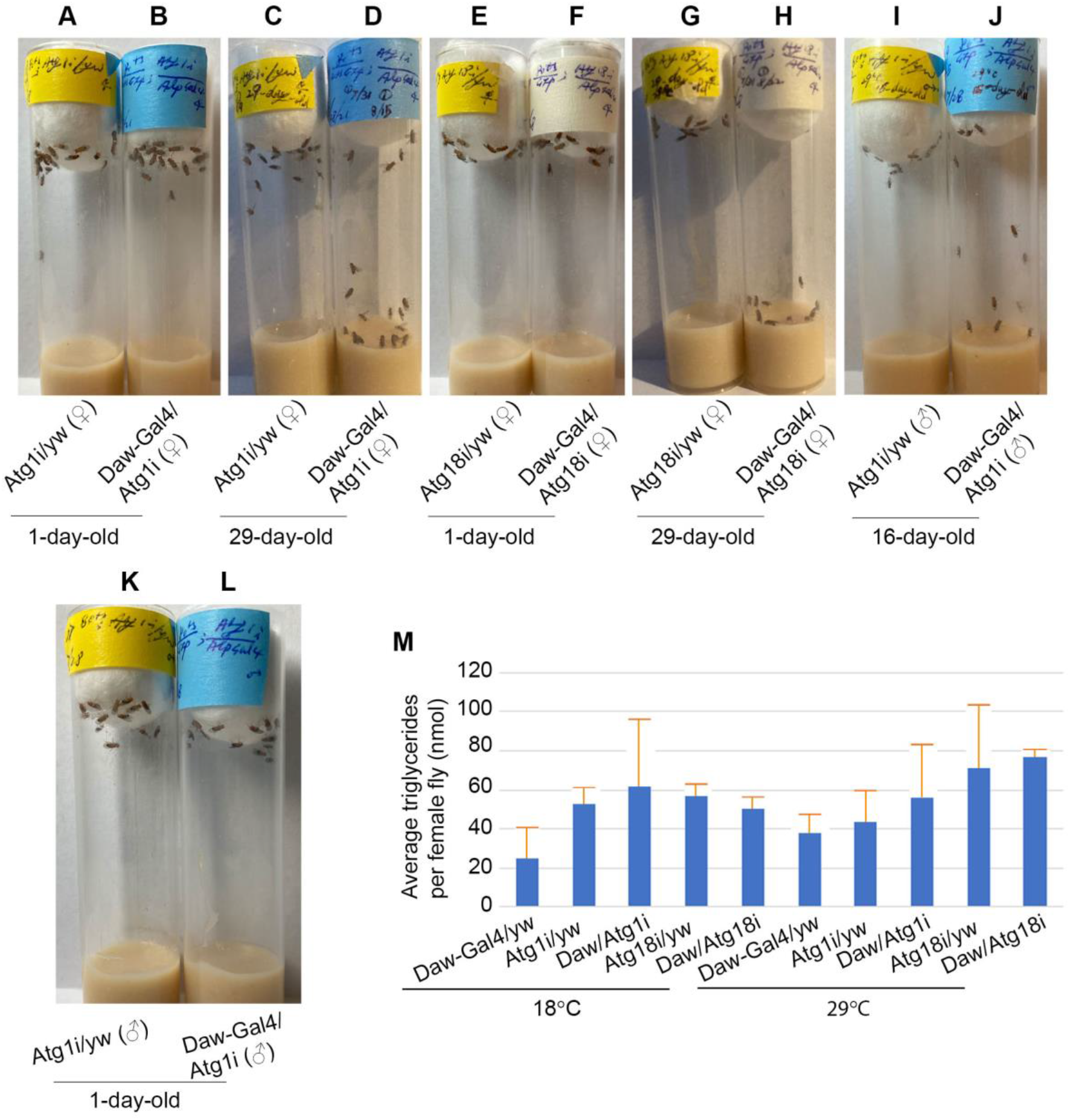
*Daw-Gal4*-induced Atg1 or Atg18 knockdown flies displayed precocious aging phenotypes (A-L) while the total lipid levels of these Atg1 or Atg18 knockdown flies are not significantly different from the controls (M). **(A, B)** Control female flies with a genotype of “*Tub-Gal80^ts^, UAS-Atg1 RNAi/yw*” (A) and experimental female flies with a genotype of “*Tub-Gal80^ts^/UAS-GFP, UAS-Atg1 RNAi/Daw-Gal4*” (B) at the age of day 1 showed no difference in their anti-gravity vial wall climbing right after they were tapped to the bottom of the vial. **(C, D)** The 29-day-old *Daw-Gal4-*induced Atg1 knockdown female flies with a genotype of “*Tub-Gal80^ts^/UAS-GFP, UAS-Atg1 RNAi/Daw-Gal4*” in vial (D) showed significantly reduced vial wall climbing speed when compared to the control flies of a genotype of “*Tub-Gal80^ts^, UAS-Atg1 RNAi/yw*” (C) of the same age and of the same sex cultured under the same temperature of 29°C (*p<0.05*). **(E, F)** Control female flies with a genotype of “*Tub-Gal80^ts^, UAS-Atg18 RNAi/yw*” (E) and the *Daw-Gal4-*induced Atg18 knockdown female flies with a genotype of “*Tub-Gal80^ts^/UAS-GFP, UAS-Atg18 RNAi/Daw-Gal4*” (F) at the age of day 1 showed no difference in their vial wall climbing after they were tapped to the bottom of the vial. **(G, H)** The 29-day-old *Daw-Gal4-*induced Atg18 knockdown female flies (H) showed delayed vial wall climbing too after these flies were tapped to the bottom of the vial as compared to the control flies in vial (G) where *Atg18 RNAi* was not expressed. **(I, J)** The significant vial wall climbing speed difference was also observed between the 16- day-old *Daw-Gal4*-induced Atg1 knockdown male flies (J) and their controls (I) (*p<0.05*). **(K, L)** Normal vial wall climbing of 1-day-old male control flies with a genotype of “*Tub-Gal80^ts^, UAS-Atg1 RNAi/yw”* (K) and young 1-day-old *Daw-Gal4-* induced Atg1 knockdown male flies with a genotype of “*Tub-Gal80^ts^/UAS-GFP, UAS- Atg1 RNAi/Daw-Gal4”* (L) showed the same vial wall climbing ability after they were tapped to the bottom of the two vials at the same time. **(M)** No statistically significant differences were observed between the triglyceride levels of the 30-day-old *Daw-Gal4-*induced Atg1RNAi or Atg18 RNAi expression female flies (*Daw/Atg1i* and *Daw/Atg18i*) and their controls (*Daw-Gal4/yw*, *Atg1i/yw*, and *Atg18i/yw*); all these flies were raised at 29°C before the assays were performed (*p>0.05*). The triglyceride levels of the 8-day-old female flies cultured at 18°C, which had the same genotypes as those raised at 29°C, were monitored for any genetic background effect on the lipid levels of these flies. No statistically significant differences of triglyceride levels were seen among these flies raised at 18°C (*p>0.05*), suggesting that the genetic backgrounds of these tested flies do not play any roles in obscuring any possible differences of triglyceride levels of the flies with Atg1 or Atg18 knocked down in adult fly adipose tissues and the control flies. Note that each orange bar of this graph represents the standard deviation from three triplicate assays of three independently prepared samples from the flies of the same genotype and of the same age. *Daw-Gal4* is also called *Alp-Gal4* as shown on the fly culture vials.

To test if the impairment of autophagy through the knockdown of the transcripts of the key autophagy genes in adipose tissue can affect the overall lipid levels of those flies, we assessed the triglyceride levels of adipose Atg1 or Atg18 knockdown flies and compared them to those from the control flies. Our experimental data showed that Atg1 or Atg18 knockdown in adipose tissue of adult flies does not affect the overall levels of triglycerides when compared to the controls (*p>0.05,* Fig. 5O).

### Over-expression of wild type Atg1 but not Atg8a gene in adult fruit fly muscle or adipose tissue shortens lifespan

Previous studies by other researchers indicated that over-expression of *Atg8a* in fly muscle extends lifespan (Bai et al., 2013), and that over-expression of wild type *Atg1* gene in fly fat body tissue can either prolong or shorten the lifespan of adult flies depending on the dosage of the protein of Atg1 (Bjedov et al., 2020). To confirm these published research data, we crossed individual compound transgenic fly lines, which included a “*Tub-Gal80^ts^, Mhc-Gal4/Tm6^TbHu^*” line and a “*CG-Gal4-UAS-GFP, Tub-Gal80^ts^*” line, to a *UAS-Atg1* or a *UAS-GFP.Atg8a* fly line or to an enhancer trap fly line in which yeast upstream activation DNA sequences (*UAS*) were inserted upstream of endogenous *Atg8a* locus on the X chromosome (*UAS-Atg8a EP*) at 18°C, and cultured their adult progeny for Atg1 or Atg8a over-expression flies together with the control flies at 29°C to induce the expression of Atg1 or Atg8a in the experimental flies. The control flies were derived from the crosses of the above parental flies for Atg1 or Atg8a over-expression to yellow white (*yw*) fruit flies.

By using the above-described experimental strategy, we found that over-expression of wild type *Atg1* gene in adult fly muscle (through *Mhc-Gal4*) or adipocytes (through *CG-Gal4*) significantly reduced the mean and maximum lifespans of those flies when compared to the controls (Fig. 6A-B, and 6C). The mean lifespans of the flies raised at 29°C with *Atg1* being over-expressed in adult fly muscle were only about 5 and 7 days as revealed with two independent *UAS-Atg1* fly lines whereas the mean lifespans of the flies from control groups of the two experiments were 27, 32, 44, and 44 days, respectively (Fig. 6A-6B). The differences between the mean lifespans of the flies with Atg1 over-expressed in adult fly muscle and the lifespans of the control flies are statistically significant in both experiments as revealed in Fig. 6A and 6B (*p <*0.05). Over-expression of wild type *Atg1* gene in adult fly adipose tissue with a *CG-Gal4* driver shortened mean lifespan by 18.5 and 21.76 days and maximum lifespan by 9 and 20 days, respectively as compared to the controls (*p*<0.05) (Fig. 6C). The specificity of wildtype *Atg1* gene in fly lifespan reduction was further confirmed by the co-expression of wildtype *Atg1* transgene and an *Atg1 RNAi* transgene in adult fly muscles (Fig. 6D). The fact that the lifespan shortening effect from over-expression of wildtype *Atg1* transgene in adult fly muscle can be effectively reversed by co-expression of wildtype *Atg1* transgene and an *Atg1 RNAi* transgene in adult fly muscles suggests that both wildtype *Atg1* transgenic line and *Atg1 RNAi* transgenic line used in our experiments are Atg1-specific fly lines, and that these fly lines are not generating any off-target effects in our experiments. All these lifespan data suggest that over-expression of wild type *Atg1* in adult fruit fly muscle or adipose tissue is detrimental to the survivorship of the tested flies.

**Figure 6.**
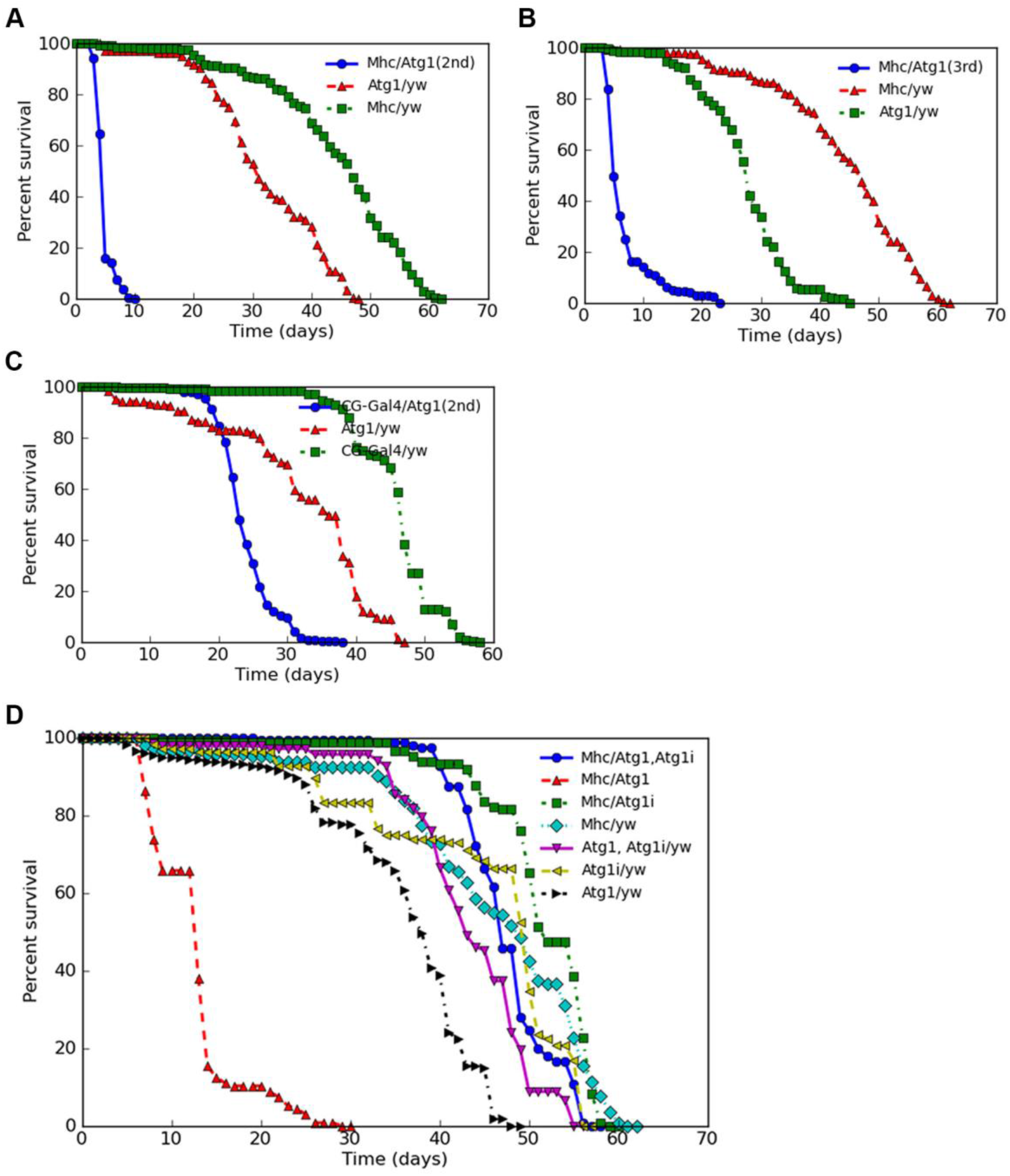
Over-expression of Atg1 in adult fly muscle or adipose tissue shortens the lifespan. **(A, B)** Over-expression of Atg1 through two independent *UAS-Atg1* fly lines with the muscle specific Gal4 driver *Mhc-Gal4* in adult male and female flies significantly reduced the mean as well as maximum lifespan when compared to the controls. The combined overall survivorships (blue curves) of Atg1 over-expression male and female flies is significantly shorter than the survivorships (red and green curves) of the control flies (*p<0.05*). **(C)** Lifespan reduction effect for both male and female flies can also be observed when wildtype *Atg1* gene was weakly expressed in the adipose tissues of adult flies through a *CG-Gal4* transgene (blue curve versus red and green curves, *p<0.05*). **(D)** The lifespan reduction effect (red curve) caused by over-expression of wildtype *Atg1* gene in adult male and female fly muscles can be abolished by co-expression of a wildtype *Atg1* gene with an *Atg1 RNAi* transgene through *Mhc-Gal4* in adult flies (blue curve). Please note that expression of *Atg1 RNAi* transgene in the adult fly muscle tissues (green curve) doesn’t shorten the lifespan of the adult male and female flies when compared to the controls (black, yellow, purple, and turquoise curves).

To determine the underlying causes of the early lethality of Atg1 over-expression flies through adult muscle, we checked if over-expression of *Atg1* gene in adult fly muscle can induce ectopic autophagy and alter the protein homeostasis and eventually cause increased levels of cell death. For autophagy test, we introduced *UAS-mCherry-Atg8a* into the *UAS-Atg1* transgenic fly line through genetic crosses and induced co-expression of mCherry-Atg8a and Atg1 in adult fly muscle through “*Tub-Gal80^ts^, Mhc-Gal4/Tm6Tm6^TbHu^*” fly line at 29°C. As demonstrated in Fig. 7A-B, over-expression of wild-type *Atg1* in adult fly muscle tissue induced ectopic autophagy as illustrated by the increased levels of mCherry signals in Fig. 7B when compared to Fig.7A. The difference of the autophagosomes induced by *Atg1* over-expression in Fig. 7B and those from the normal control muscles of Fig. 7A is about three folds. Autophagy has been implicated in helping degrade ubiquitinated protein aggregates (Ravikumar et al., 2004). Elevated levels of protein aggregates are known to affect fruit fly lifespan (Langerak et al. 2018). Ectopic expression of wild type *Atg1* in fruit fly larval fat body tissue was known to induce cellular apoptosis (Scott et al., 2004). To determine if elevated protein aggregates and cellular apoptosis are the cause of the early death of Atg1 over-expression fruit flies, we did immunostaining of 5-day-old control and Atg1 over-expression muscle tissues with an FK2 antibody, which detects mono- and polyubiquitinated proteins, and an anti-fruit fly Dcp1 antibody, which labels apoptotic cells as shown in Fig. 7G & 7H. In this immunostaining experiment, we found that over-expression of wild type *Atg1* gene in adult fly muscle caused massive increase of FK2-positive mono- and polyubiquitinated protein aggregates (Fig. 7C versus 7E, Fig. 7D versus 7F). On the other hand, apoptosis that should have been revealed by anti-Dcp1 antibody staining does not appear to be the cause for the early death of the flies with Atg1 over-expressed in the muscle (Fig. 7C-7F).

**Figure 7.**
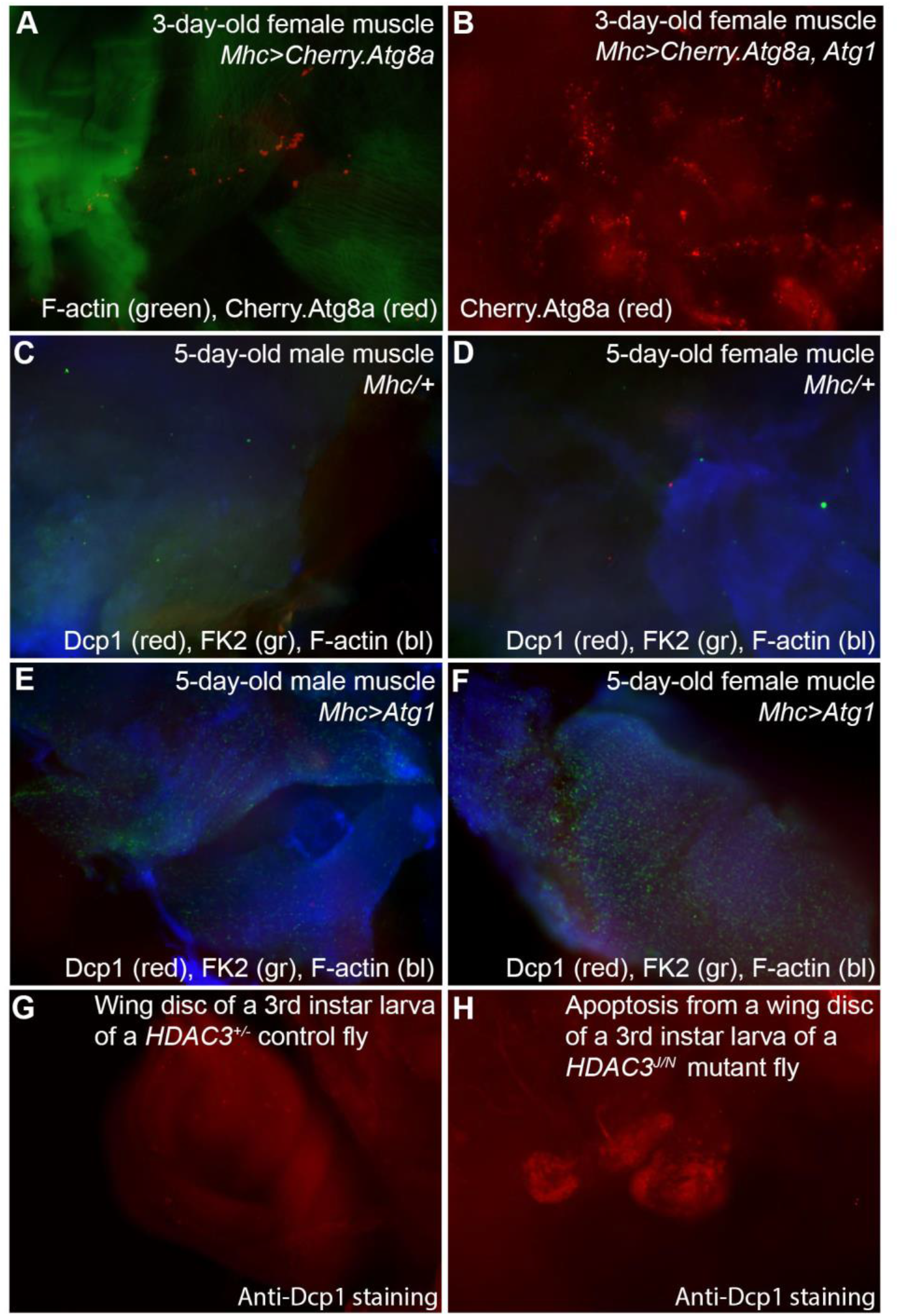
Autophagy induction by over-expression of wildtype *Atg1* gene in adult fly muscle is accompanied by accumulation of high levels of ubiquitinated protein aggregates. **(A)** Thoracic muscle of a 3-day-old female fly cultured at 29°C with a genotype of “*Tub-Gal80^ts^, Mhc-Gal4/UAS-mCherry-Atg8a*” was stained for F-actin by CytoPainter Phalloidin-iFlor 488 (green, from Abcam) and photographed for autophagosomes (red puncta) as labeled by mCherry-Atg8a fusion protein. **(B)** Ectopic autophagy (red puncta) as labeled by an mCherry-Atg8a fusion protein (red) through over-expression of wildtype *Atg1* gene in adult muscle of a 3-day-old female fly with a genotype of “*Tub-Gal80^ts^/ UAS-mCherry-Atg8a, Mhc-Gal4/UAS-Atg1*” cultured at 29°C since hatching. **(C, D)** Muscle tissues of a control male fly (C) and a control female fly (D) of 5 days of age with a genotype of “*Tub-Gal80^ts^, Mhc-Gal4/yw*”, raised at 29°C since hatching, were stained for F-actin (blue), Dcp1 (red), and ubiquitinated protein aggregates by FK2 antibody (green, Enzo Life Sciences). At this early stage of life, no apparent or very little protein aggregates (green) and apoptosis (red) can be seen from these tissues of control flies. **(E, F)** Muscle tissues of a 5-day-old male fly (E) and a 5-day-old female (F) fly with wildtype Atg1 over-expressed in their adult muscles were stained as the muscle tissues from (C) and (D) of this figure. Note that massive amount of ubiquitinated protein aggregates (green) were observed in the Atg1 over-expression muscles in (E) and (F) when compared to the control tissues in (C) and (D) of this figure. The increased levels of ubiquitinated protein aggregates from (E) and (F) do not appear to cause any increased amount of cellular apoptosis when the Dcp1 staining (red) is evaluated. **(G)** A 3^rd^ instar larval wing disc of a *HDAC3^+/-^* heterozygous control fly stained with an anti-Dcp1 antibody for apoptotic cells (red). Few sporadic apoptotic cells throughout the wing disc of this *HDAC3^+/-^* fly can be seen. **(H)** Increased levels of apoptosis are seen in the wing disc of a *HDAC3^J/N^* mutant fly were detected with the same anti-Dcp1 antibody as used for the staining of the wing disc of the control fly from (G) and for the muscle tissues as shown from (C) to (F) of this figure.

To confirm if over-expression of wildtype *Atg8a* gene in adult fly muscle extends lifespan as reported by Bai et al. (2013), we used two independent UAS-Atg8a transgenic fly lines, a *UAS-Atg8a-GFP* line and a *UAS-Atg8a* enhancer trap line (*Atg8a EP*), in our experiments. In both experiments, we could not observe any or any significant lifespan extension effect from Atg8a over-expression in adult fly muscle (*p>0.05*) (Fig. 8A-B). The 43-day mean lifespan of Atg8a over-expression flies (Mhc/GFP-Atg8a) as shown in Fig. 8A just fell between the 38-day mean lifespan and the 46-day mean lifespan of the two control groups of fruit flies. When the lifespan of the *Mhc-Gal4/UAS-Atg8aEP* fruit flies that over-expressed endogenous Atg8a in adult fly muscle was compared to those of the control flies, we found that the 47-day mean lifespan of Atg8a over-expression fruit flies at 29°C is not significantly different from the 45-day lifespan of one of two control groups of fruit flies (*p>0.05*) (Fig. 8B). All these data suggest that over-expression of Atg8a can not extend lifespan as what was previously reported. When Atg8a was over-expressed in adipose tissue through a weak *CG-Gal4* fly line, we could not see any lifespan difference of these fruit flies and their controls either (Fig. 8C).

**Figure 8.**
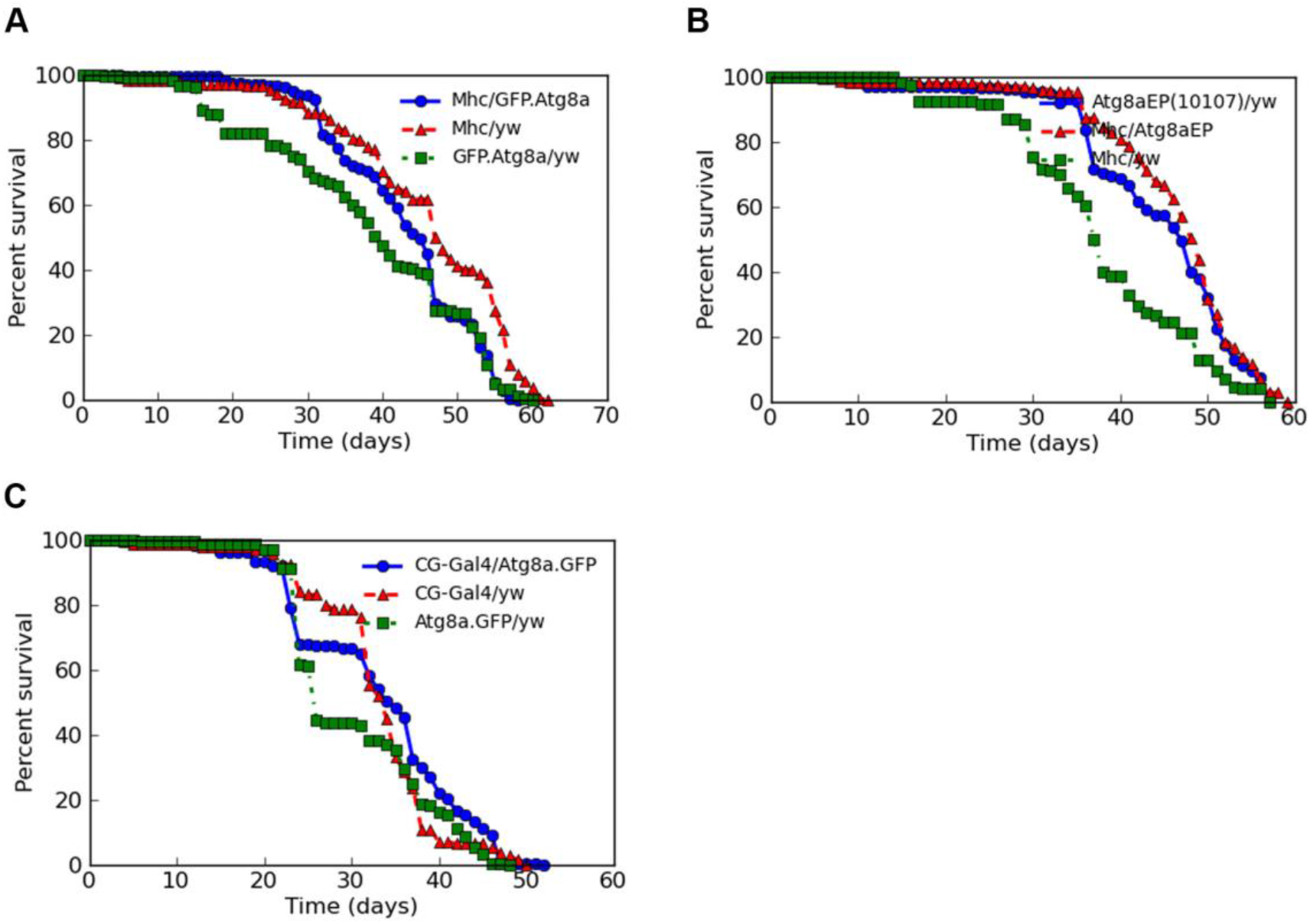
Over-expression of wildtype Atg8a in either adult fly muscle or adipose tissue does not alter the overall lifespans of the tested flies. **(A)** The lifespan of the flies with *Mhc-Gal4-*induced expression of a *UAS-GFP.Atg8a* transgene in adult fly muscle is not quite different from those lifespans of the control flies with genotypes of *Mhc/yw* or *GFP.Atg8a/yw*. **(B)** No obvious lifespan alteration effect was observed when the endogenous wildtype *Atg8a* gene on the X chromosome was over-expressed in adult muscle through an enhancer trap line, *Atg8aEP* (10107) (blue curve versus red and green curves). **(C)** Over-expression of a *UAS-GFP.Atg8a* transgene in adult adipose tissue of male and female flies through an adipose tissue-specific Gal4 driver (*CG-Gal4*) does not change the overall lifespan of these flies either when compared to those controls (blue curve versus red and green curves) (*p>0.05*).

## Discussion

Despite mounting evidence pointing to the crucial roles of autophagy genes in lifespan regulation in different animal species (Hara et al., 2006; Juhász et al., 2007; Komatsu et al., 2006; Pyo et al., 2013; and Simonsen et al., 2008), it’s unclear if these genes regulate lifespan of adult animals because of their functions in both development and adult life or just in adult life only. To delineate these possibilities, we conducted this research by using fruit fly genetics and identified the important roles of the key autophagy genes in fly lifespan regulation through adult fly adipose tissues and enteroblasts and middle gut stem cells. The lifespan regulation by At1 through enteroblasts and middle gut stem cells had previously been reported by Zhang et al. (2019), but the role of Atg1 and At18 in adipocytes for lipid metabolism, i.e. lipolysis or lipophagy, and its relationship to lifespan regulation in fruit flies as we discovered in this study are quite intriguing.

Adipose tissue works as energy reserve in flies. In fly larvae, this tissue has been identified as a target of autophagy (Scott et al., 2007). In adult fly, the same tissue has been demonstrated to be involved in fly lifespan regulation through Insulin/Tor signaling (Giannakou et al, 2004; Kapahi et al., 2004). Over-expression of Tor signaling inhibitors such as dTsc1 or dTsc2 or a dominant negative form of dS6K protein or dFoxo in fruit fly fat body tissue extends lifespan (Giannakou et al, 2004; Kapachi et al., 2004). Whether the lifespan regulation caused by the expression of Tor signaling inhibitor proteins such as dTsc1 or dTsc2 or the dominant negative form of dS6K has anything to do with altered levels of autophagy/lipophagy is not known. Although Bjedov et al. (2020) performed a study on *Atg1* in fly lifespan regulation through fly fat body tissue, the results about the effect of over-expression of *Atg1* on fly lifespan from that research are not quite clear because of the opposing effects of the Atg1 over-expression in fat body tissues on lifespan as they reported in the study. In this report, we have demonstrated that the special autophagy in fat body called lipophagy under normal fly culture conditions is very active. This active lipophagy appears to be essential for flies to achieve their normal lifespan. When lipophagy was eliminated through Atg1 or Atg18 knockdown, flies were shorter-lived and showed precocious aging phenotypes when compared to the controls. The shorter lifespans and precocious aging phenotypes of adipocyte Atg1 or Atg18 knockdown flies are not due to any obvious differences in the total lipid levels of the experimental and control flies. The shorter lifespans and precocious aging phenotypes of adipocyte Atg1 or Atg18 knockdown flies could have resulted from defective lipolysis of lipid droplets after Atg1 or Atg18 was knocked down in those cells.

Although previous published reports suggest that the key autophagy genes such as *Atg1, Atg8a,* and *Atg18* are important anti-aging regulators in adult fly muscle through over-expression of wildtype Atg8a or RNAi transgene expression for the knockdown of Atg18 (Bai et al., 2013; Xu et al., 2019), our data don’t appear to support these claims directly. These data discrepancy could have stemmed from the differences in experimental design and the Gal4 lines used in these experiments. Any future studies through more stringent loss-of-function assays such as adult tissue-specific knockouts of these genes could help reveal the important functions of the key autophagy genes in adult fly muscle and glial cells. Over-expression of Atg1 in adult fly muscle or adipocytes consistently reduced the mean as well as maximum lifespans of the tested flies whereas over-expression of wildtype Atg8a in adult fly muscle or adipocytes can’t change the overall lifespans of the tested flies at all. The lifespan reduction effect of the over-expression of Atg1 correlates with the accumulation of massive amount of ubiquitinated protein aggregates in muscle tissues. The ubiquitinated protein aggregates rather than apoptosis appeared to be the main cause of the reduced lifespan of Atg1 over-expressing flies. This data suggests that Atg1 works like a double-edged sword, which is on the one hand a potent autophagosome inducer and is required for flies to achieve their normal lifespan, but on the other hand excessive amount of Atg1 can significantly affect the normal protein homeostasis and can cut flies’ lifespan very short. Because of the dual functions of Atg1 in fly lifespan regulation, using wildtype Atg1 to prolong normal fly lifespan does not sound a viable strategy at this moment.

Autophagy can be either Atg1-dependent or Atg1-independent (Braden and Neufeld, 2013). In this study, we only assessed the function of Atg1-dependent autophagy in fly lifespan regulation and have not touched the Atg1-independent pathway that involves *Drosophila* Unc-51-like kinase (ADUK), an Atg1-independent autophagy inducer (Braden and Neufeld, 2013). For Atg1-dependent autophagy, we suggest that additional more stringent loss-of-function studies for the major autophagy genes should be performed to help unravel the potential functions of this pathway in adult fly lifespan regulation through the tissues such as adult fly muscle, neurons, and glial cells. For Atg1-independent autophagy, any tools that will target fruit fly *ADUK* gene in adult fly should suffice to provide any experimental evidence to reveal a potential function of this gene in fly lifespan regulation.

In summary, our research has uncovered an important function of the key autophagy genes such as *Atg1* and *Atg18* in adult fly lifespan regulation through adult adipocytes, enteroblasts, and gut stem cells, and at the same time we also provided evidence to show that over-expression of Atg8a in adult fly muscle or adipocytes does not seem to be able to generate any lifespan changing effects, and we cautioned the assumption that autophagy induced by over-expression of Atg1 is capable of lifespan extension in fruit flies too.

## Materials and Methods

### Fruit fly culture and genetics

All fruit fly lines used in this research were cultured on regular fruit fly food as reported before (Langerak et al., 2018). *Daw-Gal4* (on 3^rd^ chromosome) fly line was used and described previously (Langerak et al., 2018). The following fly lines used in this research were obtained from Bloomington Fly Stock Center: 1. *Tub-Gal80^ts^* (on 2^nd^ chromosome); 2. *Tub-Gal80^ts^* (on 3^rd^ chromosome); 3. *Mhc-Gal4/Tm3^Sb^*; 3. *CG-Gal4* (on 2^nd^ chromosome); 4. *UAS-mCherry-Atg8a* (on 2^nd^ chromosome, stock 37749, Chang and Neufeld, 2009); 5. *UAS-Atg1* (on 2nd chromosome, stock 51654); 6. *UAS- Atg1* (on 3^rd^ chromosome, stock 51655); 7. *UAS-Atg1 RNAi* (on 2^nd^ chromosome, stock 44034); 8. *UAS-Atg1 RNAi* (on 3^rd^ chromosome, stock 26731); 9. *UAS-Atg1 RNAi* (on 3^rd^ chromosome, stock 35177); 10. *UAS-Atg5 RNAi* (on 3^rd^ chromosome, stock 27551); 11. *UAS-Atg5 RNAi* (on 3^rd^ chromosome, stock 34899); 12. *EP-Atg8a* (on X chromosome, stock 10107); 13. *UAS-Atg9 RNAi* (on 3^rd^ chromosome, Stock 34901); 14. *UAS-Atg18 RNAi* (on 3^rd^ chromosome, stock 28061); 15. *Esg-Gal4*; and 16*. Repo-Gal4/Tm3^Sb^*. The TRiP RNAi fly lines obtained from Bloomington Fruit Fly Stock Center were created in *y^[1]^v^[1]^* genetic background and deposited to the center by Perkins et al. (2015). The RNA knockdown effect of the *UAS-Atg9 RNAi* fly line (Stock 34901) and *UAS-Atg18 RNAi* (Stock 2806) fly line from Bloomington Stock Center had been demonstrated through qPCR by Xu et al. (2019) and in this article as illustrated in Fig.1G-1I & 1J. All the RNAi fly lines of autophagy genes used in this research are in y^[1]^v^[1]^ genetic background. The following compound transgenic fly lines were created by using different second and third balancer chromosomes through fly mating and genetics: 1. “*Tub-Gal80^ts^, Tub-Gal4/Tm6^TbHu^”*; 2. “*Tub-Gal80^ts^, Mhc-Gal4/Tm6^TbHu^”*; 3. “*CG-Gal4-UAS-GFP, Tub-Gal80^ts^*”; 4. “*Tub-Gal80^ts^, UAS-Atg1 RNAi”*; 5. “*Tub-Gal80^ts^, UAS- Atg18 RNAi”*; 6. “*UAS-Atg1, UAS-Atg1RNAi*”; 7. “*UAS-Atg1, UAS-Atg9 RNAi*”; and 8. “*UAS-Atg1, UAS-Atg18 RNAi*”. Most of our Gal4 lines are in a *w^[1118]^* genetic background. *HDAC3^J^/Tm6^TbHu^* and *HDAC3^N^/Tm6^TbHu^* mutant fruit flies used in this study were published before (Zhu et al., 2008).

### Lifespan and starvation resistance assays

All the lifespan analyses for Atg knockdown or over-expression flies throughout this research project were carried out either by crossing individual *Tub-Gal80^ts^* associated *Gal4* lines to individual *UAS-Atg1 RNAi*, *UAS-Atg5 RNAi*, *UAS-Atg9 RNAi*, *UAS-Atg18 RNAi, UAS-Atg1,* or *UAS-Atg8a* fly lines, or by mating different *Tub-Gal80^ts^* associated *UAS-Atg1 RNAi* or *UAS-Atg18 RNAi* fly lines to individual *Gal4* lines such as *Repo-Gal4* or *Daw-Gal4* at 18°C, and culturing their newly hatched progenies that inherited the corresponding transgenes at 29°C. All the control flies, which were derived from the newly hatched progenies from the mating of the parental flies used for the Atg knockdown or ever-expression to a yellow white fly line (*yw*) at 18°C, were sorted at room temperature and cultured under the same 29°C just as how the Atg knockdown or over-expression flies were sorted and cultured. We did not use RU- 486 based gene switch transgenic fly lines such as *Da-GS* or *Tub-GS* in our lifespan study just because these fly lines can induce the expression of any transgene that’s under the UAS control even without the treatment of the flies with RU-486 as we demonstrated before (Langerak et al., 2018). The leaky induction of transgene expression by *Da-GS* or *Tub-GS* could cause confounding effects for fly lifespan analysis. In our experiments, about 150 to 200 newly hatched flies of each genotype with males and females being at about equal representation were sorted and cultured in different vials for lifespan study. Each culture vial hosted on average 15 to 25 male or female flies. We only compared the lifespans of different genotypes of flies from a single experiment performed at the same time and never compared the lifespans of flies from different experiments conducted at different times because any subtle differences of fly culture conditions from different experiments performed at different times can change the lifespans of tested flies easily. For lifespan data recording, dead flies from each culture vial were inspected and recorded either daily or every other day. Living flies from individual old vials were transferred to new vials every two to three days to ensure the freshness of the food. The 12-hour light and dark cycle was maintained throughout the fly culture. The lifespan data of all the experiments were analyzed by an online lifespan analysis tool called “Oasis” (Yang et al., 2011). Both Log-Rank and Fisher’s Exact statistical methods were used for computing the P-values for lifespan differences between experimental and control flies. For starvation resistance analysis, female or male fruit flies of different genotypes were placed in the fly culture vials containing water-soaked cotton balls at either 18°C or 29°C. Each vial hosted about 15 to 20 flies. Each genotype of flies of each sex was tested in three independent vials for the survivorship under the condition of starvation. Dead flies were recorded at specific time points. The whole starvation experiment was deemed to be done when no surviving flies were seen anymore.

### Imaging of mCherry-Atg8a-labeld autophagosomes

The live thoracic indirect flight muscles of control, Atg1 knockdown, and Atg18 knockdown flies with the expression of mCherry-Atg8a fusion protein were counterstained with diluted CytoPainter Phalloidin-iFlor 488 (from Abcam) for F-actin at room temperature for 15 minutes before the tissues were mounted in phosphate-buffered saline solution (PBS) on slides for imaging. Newly dissected brains of control, Atg1 knockdown, and Atg18 knockdown flies expressing mCherry-Atg8a fusion protein were mounted in PBS on slides before pictures were taken. Live adipocytes of control, Atg1 knockdown, and Atg18 knockdown flies expressing mCherry-Atg8a fusion protein were stained with 4’,6-diamidino-2-phenylindole (DAPI) for nuclei for 10 minutes before they were mounted in PBS on slides and photographed. All the fluorescent pictures were taken with a Nikon fluorescence compound microscope equipped with a Teledyme Qimaging Micropublishing 6 digital camera.

### Quantitative polymerase chain reaction (qPCR)

Each RNA sample was extracted and precipitated from the homogenized tissues of two adult flies (one female and one male) in Trizol according to the manufacturer’s instructions (Zymo Research R2060). For each genotype of fruit flies, three independent individual RNA samples were prepared and used for three independent qPCR reactions. For qPCR, the extracted RNA was quantified using a nanodrop, and 1.2 ng RNA was used as a template for reverse transcription. cDNA was synthesized by using SuperScript III (Invitrogen 18080093). qPCR was performed by using SYBR Green reagent (Roche 04707516001) on a Light Cycler 480 according to the manufacturer’s instructions. The transcripts of the housekeeping gene *Rpl23* were used for normalization. The two primers of *Atg 1* gene used in this experiment are 5’-CTAAAGCCGTCGTCCAATGT-3’ (forward) and 5’- GAACAGCATGCTCCGGTATT-3’ (reverse). Primers of the *Rpl23* gene used: 5’- GTTTGCGCTGCCGAATAACCAC-3’ (forward) and 5’-GACAACACCGGAGCCAAGAACC-3’(reverse).

### Immunostaining and imaging

Immunostaining of adult fly tissues was performed as described previously (Langerak et al., 2018). The following antibodies and reagents were purchased and used in this research: Rabbit anti-cleaved *Drosophila* Dcp-1 (Asp216) antibody from Cell Signaling Technology, Inc. (1:100 dilution was used to monitor cellular apoptosis, catalog # 9578S); Mouse monoclonal antibody anti-Mono- and poly-ubiquitinylated proteins (FK2) from Enzo Life Sciences (1:200 dilution was used), Alexa Fluor 488 anti-mouse IgG (H+L) (Catalog # 4408S) and Alexa Fluor 488 anti-rabbit IgG (H+L) from Cell Signaling Technology (1:200 dilution was used), CytoPainter phalloidin-iFluor 488 reagent (catalog # ab176753) from Abcam, and Phalloidin-iFluor 350 reagent (catalog # ab176751) from Abcam. All the images were taken with a Nikon compound fluorescence microscope.

### Triglyceride assay

After the removal of the wings, every 3 to 7 female or male fruit flies were ground thoroughly at room temperature in 5% triton X-100 solution (100 µl per fly) in a mortar with a pestle. 5% triton X-100 solution was prepared by diluting Triton X-100 purchased from Sigma in distilled water. Three independent preparations of the samples of each genotype of female or male fruit flies of defined age were used in this assay. The ground tissues and the dissolved triglycerides in triton X-100 solution were collected into 1.5 ml microcentrifuge tubes and were heated in a water bath for about 2 minutes at about 80°C to 90°C before they were centrifuged at a speed of 12,000 round per minute at room temperature for two minutes to get rid of tissue debris. The supernatant containing the total triglycerides was saved at −80°C for analysis. The total triglycerides were quantified with a triglyceride assay kit (product # ab65336) from Abcam by following the standard protocol offered by the company. The whole assay was performed in a Nunc-Immuno MicroWell 96-well clear-flat-bottom plate (Thermo Fisher Scientific, Rockford, IL). The optical density at 570 nm was measured using Cytation 3 Multi-Mode Reader (Bio-Tek Instruments, Inc., Winooski, VT).

### Fly climbing assay

For aging phenotype, we tapped the control and experimental flies of the same age and of the same sex raised under the same culture conditions in fresh food vials to the bottom of the vials at the same time and let the flies climb up on the wall right after the tapping and record the differences of wall climbing speed of experimental and control flies by a smart phone camera. The faster the flies climbed to the top of the vial, the younger and healthier they should be. The longer the flies take to reach the top of the vial, the more aged they may represent. For each experiment, three independent tests were performed for each genotype of flies, and the data were analyzed through a Chi-square test statistic method for p values.

### Statistics

The lifespan data of the fruit flies of different genotypes were analyzed and compiled by using the online Oasis lifespan analysis tool where both Log-rank and Fisher’s exact statistical methods were used (Yang et al., 2011). The other statistical analyses were carried out by using either Student T-test or an ANOVA data analysis tool pack from Microsoft Excel software or a Chi-square test statistic method for the corresponding *p* values.

## Acknowledgement

We thank Mrs. Patricia Bunce, Dr. Sky Pike, Dr. Karen Barkel, Dr. Bradley Isler, and Dr. Mary Beth Zimmer for their strong support of this research project. We are grateful to Dr. Paul Klatt and Dr. Joseph Lipar for their critical reading of the manuscript. Crossroads Charter Academy students Jiahong Zhou and Owen PiiPPo volunteered in our lab during the late phase of this project. This research project has been funded by Ferris State University Ferris Foundation Exceptional Merit Grants and Ferris State University Faculty Research Grants to C.C.Z., Ferris State University Undergraduate Student Research Grants from the College of Arts, Sciences, and Education to J.C., T.P.B, D.P., B.W., and G.A., and a Ferris Student Summer Research Grant to J.Z..

## Author contributions

Conceptualization: C.C.Z.; Data Curation: C.C.Z.; Formal Analysis: C.C.Z.; Funding Acquisition: C.C.Z.; Investigation: M.B., J.C., T.P.B., H.B, M. R, J.Z., D.P., L.W., B.W., J.V.W., B.W., G.A., Z.H., E.D., A.A., E.K., F.A., C.C.Z.; Methodology: C.C.Z.; Project Administration: C.C.Z.; Resources: C.C.Z., M.B.O., T.P.N.; Software: C.C.Z.; Supervision: C.C.Z.; Validation: C.C.Z; Visualization: C.C.Z.; Writing – original draft: C.C.Z; Writing – review & editing: T.P.N, C.C.Z..

**The authors declare that they have no conflict of interest.**

---

**Supplemental Figure 1.**
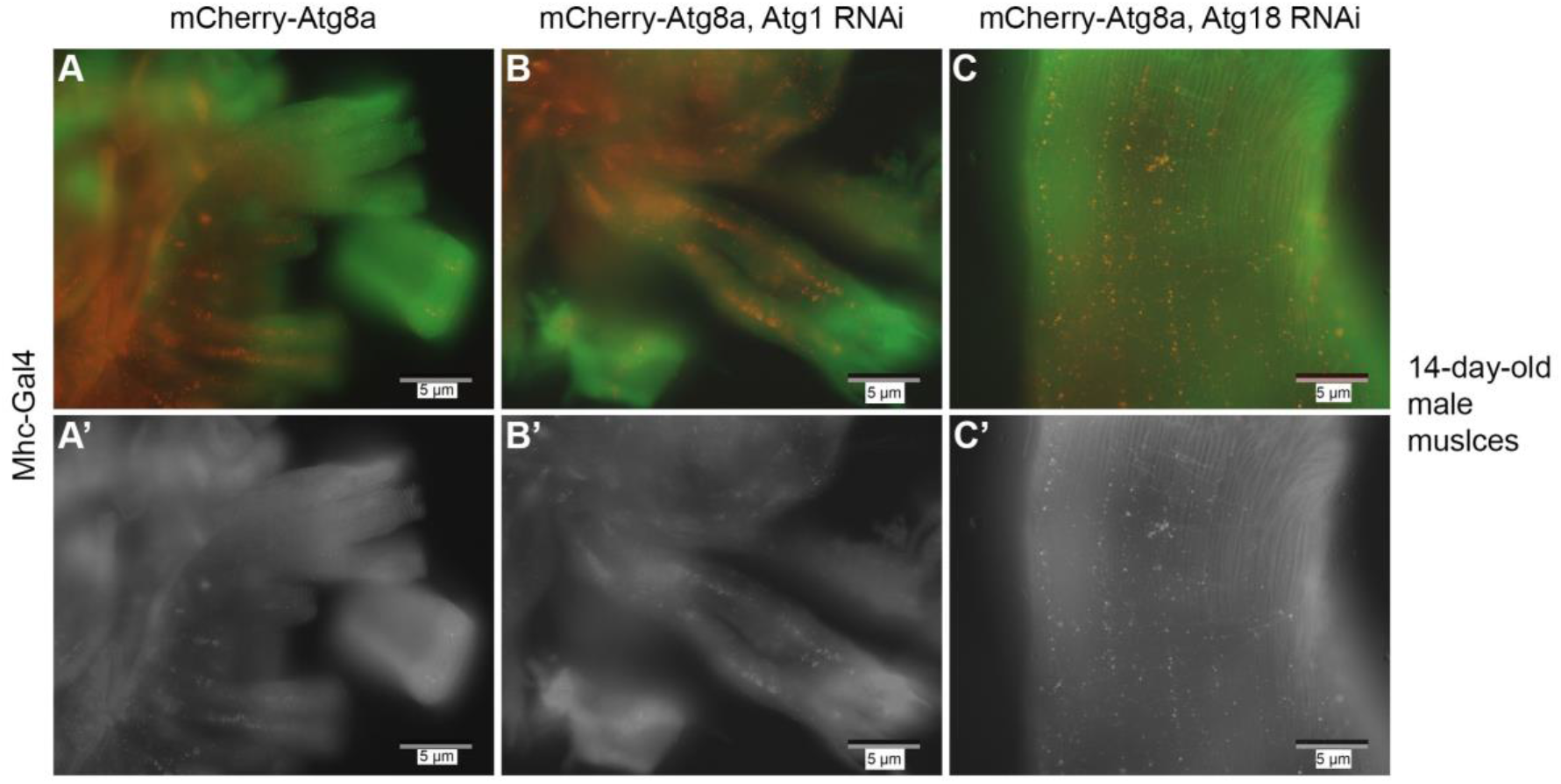
M*h*c*-Gal4*-induced expression of *Atg1 RNAi* or *Atg18 RNAi* in adult male fly muscles does not change the overall levels of mCherry-Atg8a-labeled autophagosomes. **(A-C)** mCherry-Atg8a-labeled autophagosomes (red puncta) can well be observed in CytoPainter Phalloidin-iFlor 488 (green)- marked live thoracic indirect flight muscles of three 14-day-old male flies with genotypes of “*Tub-Gal80^ts^/UAS-mCherry-Atg8a, Mhc-Gal4/+*” (A), “*Tub-Gal80^ts^/UAS- mCherry-Atg8a, Mhc-Gal4/UAS-Atg1 RNAi*” (B), and “*Tub-Gal80^ts^/UAS-mCherry-Atg8a, Mhc-Gal4/UAS-Atg18 RNAi*” (C), respectively. **(A’-C’)** Gray images of (A-C) of this figure.

**Supplemental table 1.**
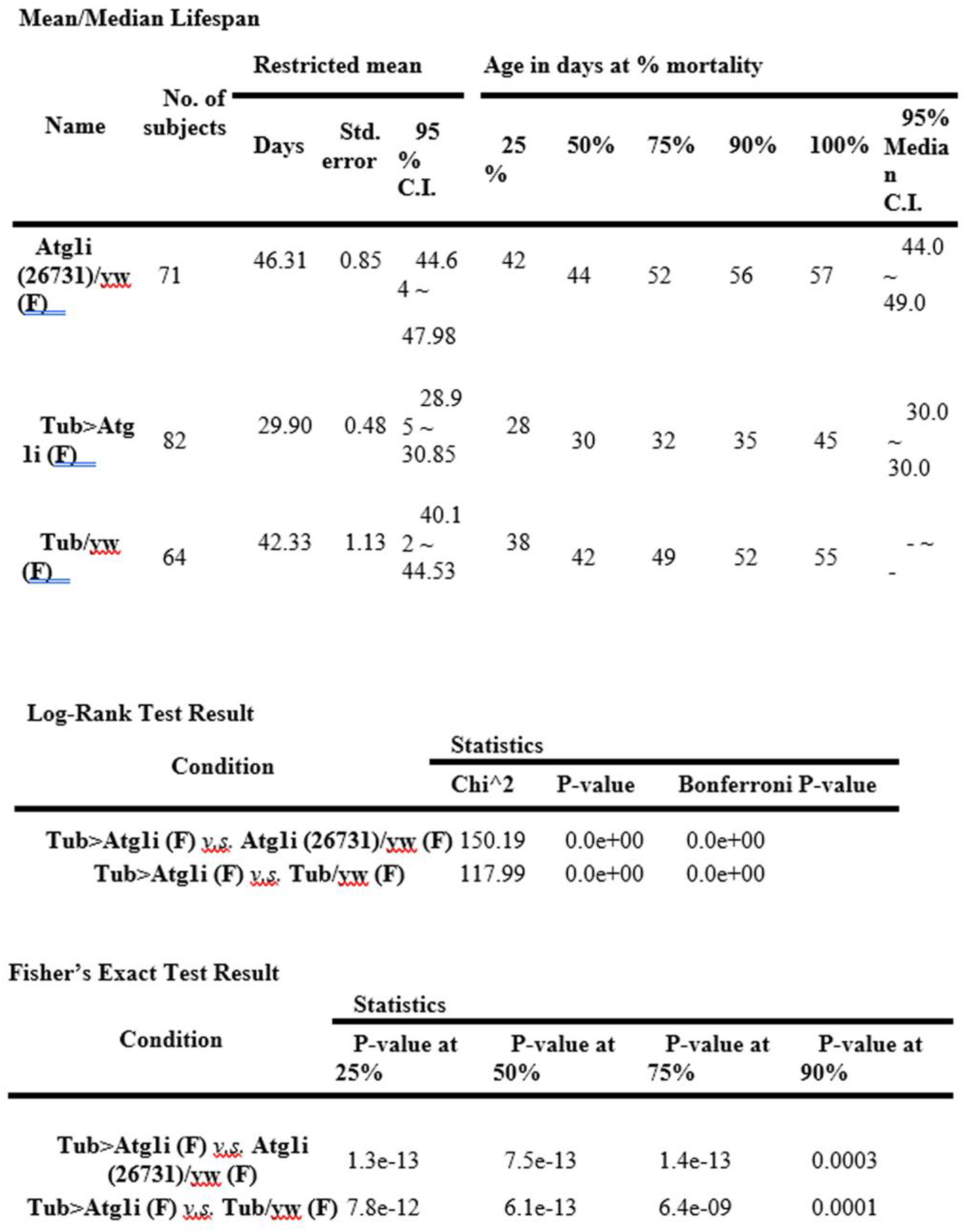
Statistical data on the lifespans of ubiquitous adult-specific Atg1 knockdown female (Tub>Atg1i (F)) and control female files (Atg li(26731)/yw (F) and Tub/yw (F))

**Supplemental table 2.**
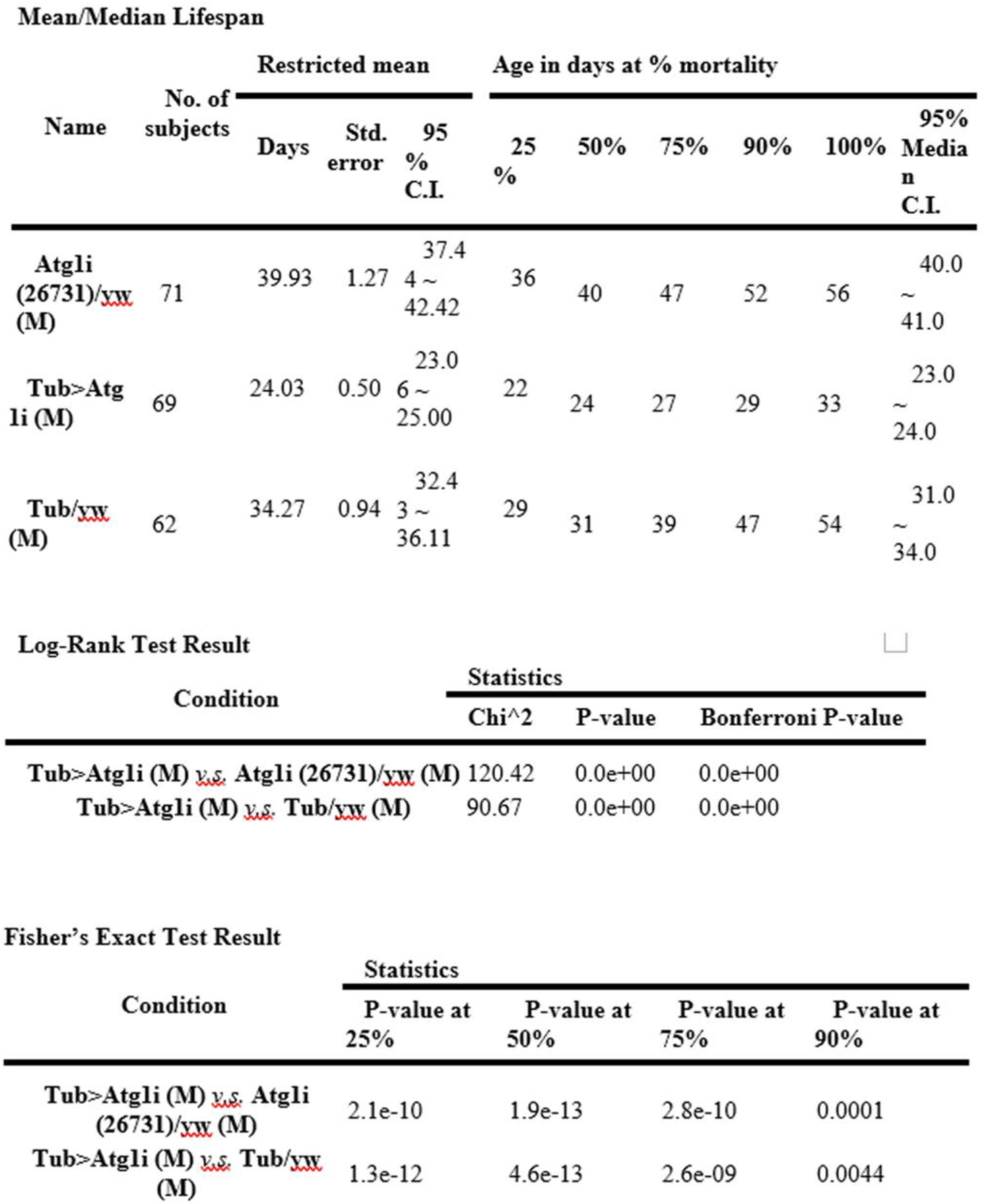
Statistical data on the lifespans of ubiquitous adult-specific Atg1 knockdown male (Tub>Atg1i (M)) and control female files (Atg li(26731)/yw (M) and Tub/yw (M))

**Supplemental table 3.** Statistical data on the lifespans of *Tub-Gal4* induced adult Atg9 knockdown male and female flies (*TubGal4/Atg9i*) and their controls (*UAS-Atg9i(34901)/yw* and *TubGal4/yw*) reared at 29°C

Mean/Median Lifespan
| Name | No. of subjects | Restricted mean |  |  | Age in days at % mortality |  |  |  |  |  |
| --- | --- | --- | --- | --- | --- | --- | --- | --- | --- | --- |
|  |  | Days | Std. error | 95% C.I. | 25 % | 50% | 75% | 90% | 100% | 95% Median C.I. |
| UAS-Atg9i (34901)/yw | 175 | 39.02 | 0.65 | 37.7<br>5 ~<br>40.28 | 32 | 40 | 46 | 49 | 55 | 38.0<br>~<br>41.0 |
| TubGal4/Atg9i | 187 | 25.70 | 0.61 | 24.5<br>0 ~<br>26.89 | 21 | 25 | 33 | 37 | 40 | 25.0<br>~<br>27.0 |
| TubGal4/yw | 163 | 36.64 | 0.73 | 35.2<br>1 ~<br>38.08 | 30 | 34 | 43 | 50 | 60 | 32.0<br>~<br>36.0 |

| Condition | Statistics |  |  |  |
| --- | --- | --- | --- | --- |
|  | P-value at 25% | P-value at 50% | P-value at 75% | P-value at 90% |
| *TubGal4/Atg9i v.s. UAS-Atg9i(34901)/yw* | 9.0e-13 | 2.2e-12 | 9.7e-13 | 2.2e-09 |
| *TubGal4/Atg9i v.s. TubGal4/yw* | 7.5e-13 | 8.2e-13 | 1.6e-12 | 2.3e-11 |

| Condition | Statistics |  |  |
| --- | --- | --- | --- |
|  | Chi^2 | P-value | Bonferroni P-value |
| *TubGal4/Atg9i v.s. UAS-Atg9i(34901)/yw* | 187.23 | 0.0e+00 | 0.0e+00 |
| *TubGal4/Atg9i v.s. TubGal4/yw* | 106.53 | 0.0e+00 | 0.0e+00 |

**Supplemental table 4.** Statistical data on the combined lifespans of *Tub-Gal4* induced adult Atg18 knockdown male and female flies (*TubGal4/Atg18i*) and their controls (*Atg18i(28061)/yw* and *TubGal4/yw*) reared at 29°C

Mean/Median Lifespan
| Name | No. of subjects | Restricted mean |  |  | Age in days at % mortality |  |  |  |  |  |
| --- | --- | --- | --- | --- | --- | --- | --- | --- | --- | --- |
|  |  | Days | Std. error | 95% C.I. | 25 % | 50% | 75% | 90% | 100% | 95% Median C.I. |
| *Atg18i(28061)/yw* | 199 | 41.16 | 0.47 | 40.23 ~ 42.09 | 37 | 42 | 46 | 49 | 52 | 42.0 ~ 43.0 |
| *Tub-Gal4/Atg18i* | 204 | 23.64 | 0.22 | 23.20 ~ 24.07 | 22 | 24 | 26 | 28 | 31 | 23.0 ~ 23.0 |
| *Tub-Gal4/yw* | 191 | 37.89 | 0.64 | 36.64 ~ 39.14 | 30 | 40 | 45 | 49 | 53 | 37.0 ~ 40.0 |

| Condition | Statistics |  |  |  |
| --- | --- | --- | --- | --- |
|  | P-value at 25% | P-value at 50% | P-value at 75% | P-value at 90% |
| *Tub-Gal4/Atg18i* v.s. *Atg18i(28061)/yw* | 1.3e-12 | 9.0e-13 | 8.0e-13 | 5.4e-13 |
| *Tub-Gal4/Atg18i* v.s. *Tub-Gal4/yw* | 4.9e-13 | 1.7e-12 | 1.3e-12 | 2.9e-13 |

| Condition | Statistics |  |  |
| --- | --- | --- | --- |
|  | Chi^2 | P-value | Bonferroni P-value |
| *Tub-Gal4/Atg18i* v.s. *Atg18i(28061)/yw* | 428.61 | 0.0e+00 | 0.0e+00 |
| *Tub-Gal4/Atg18i* v.s. *Tub-Gal4/yw* | 293.46 | 0.0e+00 | 0.0e+00 |

**Supplemental table 5.**
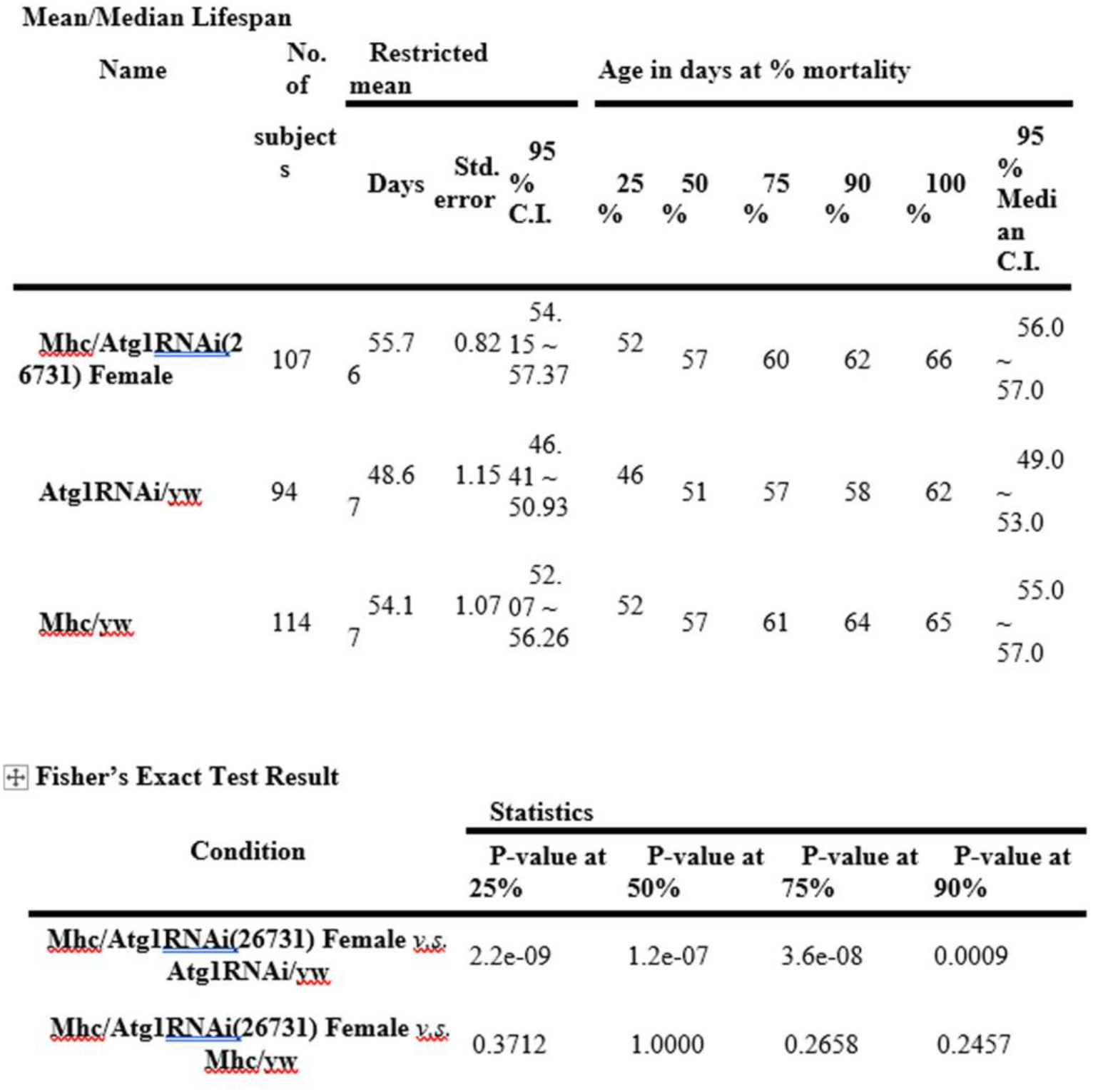
Statistical data on the lifespans of adult muscle-specific Atg1 knockdown females flies (Mhc/Atg1RNAi(26731)) and control female flies (Atg1RNAi/yw, Mhc/yw)

**Supplemental table 6.** Statistical data on the lifespans of male Atg1 muscle knockdown files (Mhc/Atg1i (26731)) and their controls

Mean/Median Lifespan
| Name | No. of subjects | Restricted mean |  |  | Age in days at % mortality |  |  |  |  |  |
| --- | --- | --- | --- | --- | --- | --- | --- | --- | --- | --- |
|  |  | Days | Std. error | 95 % C.I. | 25 % | 50% | 75% | 90% | 100% | 95% Median C.I. |
| <u>Mhc/Atgli (26731) (M)</u> | 107 | 45.93 | 0.67 | 44.6<br>3 ~ 47.24 | 44 | 47 | 51 | 54 | 56 | 45.0<br>~ 47.0 |
| <u>Atgli/yw (M)</u> | 114 | 44.79 | 1.12 | 42.5<br>9 ~ 46.99 | 42 | 49 | 53 | 56 | 57 | 47.0<br>~ 49.0 |
| <u>Mhc/yw (M)</u> | 106 | 41.06 | 0.71 | 39.6<br>7 ~ 42.45 | 36 | 41 | 46 | 51 | 58 | 39.0<br>~ 42.0 |

Log-Rank Test Result
| Condition | Statistics |  |  |  |
| --- | --- | --- | --- | --- |
|  | P-value at 25% | P-value at 50% | P-value at 75% | P-value at 90% |
| <u>Mhc/Atgli (26731) (M)</u> <u>vs.</u> <u>Atgli/yw (M)</u> | 0.4511 | 0.1763 | 0.0797 | 0.0060 |
| <u>Mhc/Atgli (26731) (M)</u> <u>vs.</u> <u>Mhc/yw (M)</u> | 3.1e-05 | 3.1e-05 | 0.0004 | 0.2398 |

**Supplemental table 7.** Statistical data on the combined lifespans of *Esg-Gal4* induced adult Atg1 knockdown male and female flies (*Esg-Gal4/Atg1i*(26731)) and their controls (*Atg1i/yw* and *Esg-Gal4/yw*) cultured at 29°C

Mean/Median Lifespan
| Name | No. of<br>subject<br>s | Restricted mean |  |  | Age in days at % mortality |  |  |  |  |  |
| --- | --- | --- | --- | --- | --- | --- | --- | --- | --- | --- |
|  |  | Days | Std.<br>error | 95<br>%<br>C.I. | 25<br>% | 50% | 75% | 90% | 100% | 95<br>%<br>Medi<br>an<br>C.I. |
| Esg-Gal4/Atg1i(26731) | 150 | 32.42 | 0.90 | 30.66 ~ 34.18 | 29 | 35 | 40 | 44 | 49 | 33.0 ~ 36.0 |
| Atg1i/yw | 165 | 40.82 | 0.73 | 39.40 ~ 42.25 | 36 | 43 | 46 | 51 | 56 | 41.0 ~ 43.0 |
| Esg-Gal4/yw | 148 | 38.76 | 0.77 | 37.25 ~ 40.26 | 34 | 41 | 45 | 48 | 54 | 38.0 ~ 41.0 |

| Condition | Statistics |  |  |  |
| --- | --- | --- | --- | --- |
|  | P-value at 25% | P-value at 50% | P-value at 75% | P-value at 90% |
| **Esg-Gal4/Atg1i(26731) v.s. Atg1i/yw** | 5.6e-07 | 6.8e-09 | 8.0e-13 | 8.8e-06 |
| **Esg-Gal4/Atg1i(26731) v.s. Esg-Gal4/yw** | 7.9e-06 | 2.5e-05 | 1.5e-06 | 4.9e-07 |

**Supplemental table 8.**
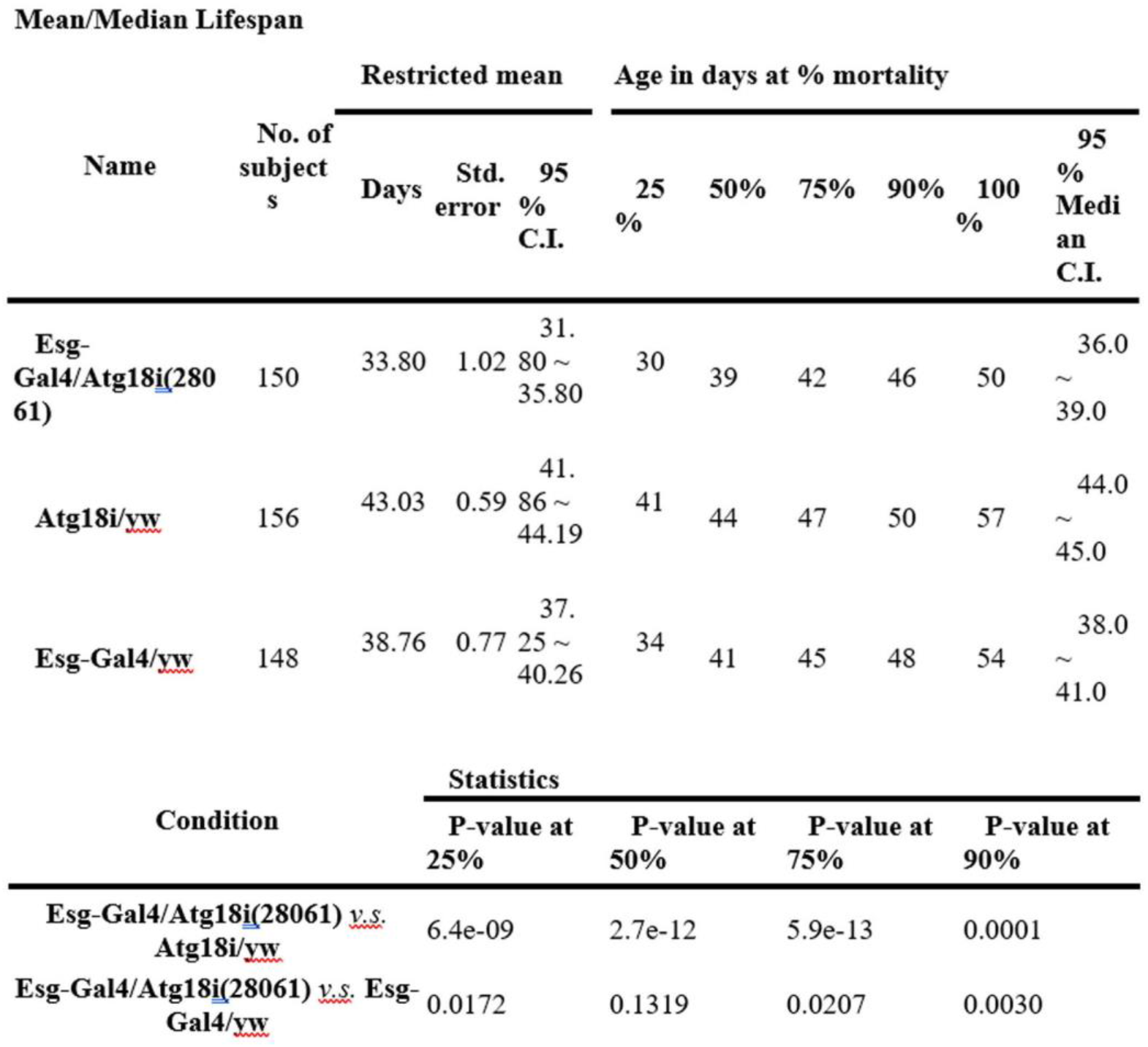
Statistical data on the combined lifespans of *Esg-Gal4* induced adult Atg18 knockdown male and female flies (*Esg-Gal4/Atg18i*(28061)) and their controls (*Atg18i/yw* and *Esg-Gal4/yw*) cultured at 29°C

Mean/Median Lifespan
| Name | No. of subjects | Restricted mean |  |  | Age in days at % mortality |  |  |  |  |  |
| --- | --- | --- | --- | --- | --- | --- | --- | --- | --- | --- |
|  |  | Days | Std. error | 95 % C.I. | 25 % | 50% | 75% | 90% | 100 % | 95 % Median C.I. |
| Esg-Gal4/Atg18i(28061) | 150 | 33.80 | 1.02 | 31.80 ~ 35.80 | 30 | 39 | 42 | 46 | 50 | 36.0 ~ 39.0 |
| Atg18i/yw | 156 | 43.03 | 0.59 | 41.86 ~ 44.19 | 41 | 44 | 47 | 50 | 57 | 44.0 ~ 45.0 |
| Esg-Gal4/yw | 148 | 38.76 | 0.77 | 37.25 ~ 40.26 | 34 | 41 | 45 | 48 | 54 | 38.0 ~ 41.0 |

| Condition | Statistics |  |  |  |
| --- | --- | --- | --- | --- |
|  | P-value at 25% | P-value at 50% | P-value at 75% | P-value at 90% |
| **Esg-Gal4/Atg18i(28061) v.s. Atg18i/yw** | 6.4e-09 | 2.7e-12 | 5.9e-13 | 0.0001 |
| **Esg-Gal4/Atg18i(28061) v.s. Esg-Gal4/yw** | 0.0172 | 0.1319 | 0.0207 | 0.0030 |

**Supplemental table 9.**
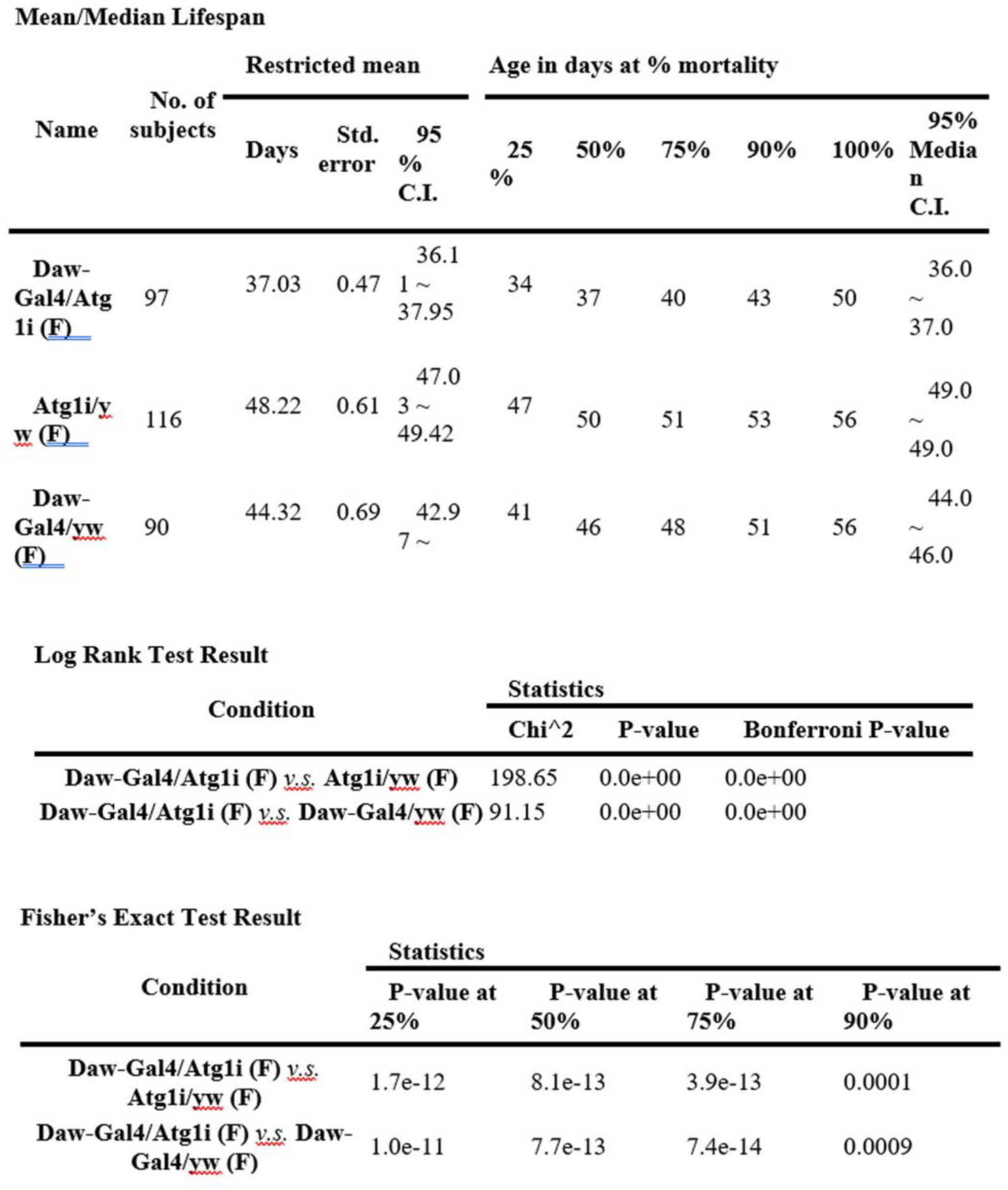
Statistical data on the lifespans of *Daw-Gal4* induced adult Atg1 knockdown female flies (*Daw Gal4/Atg1i (F)*) and their controls (*Atg1i/yw (F)* and *Daw-Gal4/yw (F)*) cultured at 29°C

**Supplemental table 10.** Statistical data on the lifespans of *Daw-Gal4* induced adult Atg1 knockdown male flies (*Daw Gal4/Atg1i (M)*) and their controls (*Atg1i/yw (M)* and *Daw-Gal4/yw (M)*) cultured at 29°C

Mean/Median Lifespan
| Name | No. of subjects | Restricted mean |  |  | Age in days at % mortality |  |  |  |  |  |
| --- | --- | --- | --- | --- | --- | --- | --- | --- | --- | --- |
|  |  | Days | Std. error | 95% C.I. | 25% | 50% | 75% | 90% | 100% | 95% Median C.I. |
| **Daw-Gal4/Atg1i** (M) | 89 | 23.15 | 0.58 | 22.0<br>1 ~ 24.28 | 19 | 24 | 26 | 29 | 41 | 23.0<br>~ 24.0 |
| **Atg1i/yw** (M) | 106 | 37.01 | 0.61 | 35.8<br>0 ~ 38.21 | 34 | 35 | 42 | 46 | 49 | 34.0<br>~ 36.0 |
| **Daw-Gal4/yw** (M) | 101 | 32.36 | 0.80 | 30.7<br>9 ~ 33.92 | 27 | 33 | 37 | 43 | 47 | 32.0<br>~ 34.0 |

Log Rank Test Result
| Condition | Statistics |  |  |
| --- | --- | --- | --- |
|  | Chi^2 | P-value | Bonferroni P-value |
| **Daw-Gal4/Atg1i** (M) *v.s.* **Atg1i/yw** (M) | 185.37 | 0.0e+00 | 0.0e+00 |
| **Daw-Gal4/Atg1i** (M) *v.s.* **Daw-Gal4/yw** (M) | 86.11 | 0.0e+00 | 0.0e+00 |

Fisher's Exact Test Result
| Condition | Statistics |  |  |  |
| --- | --- | --- | --- | --- |
|  | P-value at 25% | P-value at 50% | P-value at 75% | P-value at 90% |
| **Daw-Gal4/Atg1i** (M) *v.s.* **Atg1i/yw** (M) | 3.7e-13 | 5.8e-13 | 2.3e-12 | 7.2e-06 |
| **Daw-Gal4/Atg1i** (M) *v.s.* **Daw-Gal4/yw** (M) | 1.9e-07 | 1.0e-12 | 2.8e-12 | 0.0004 |

**Supplemental table 11.**
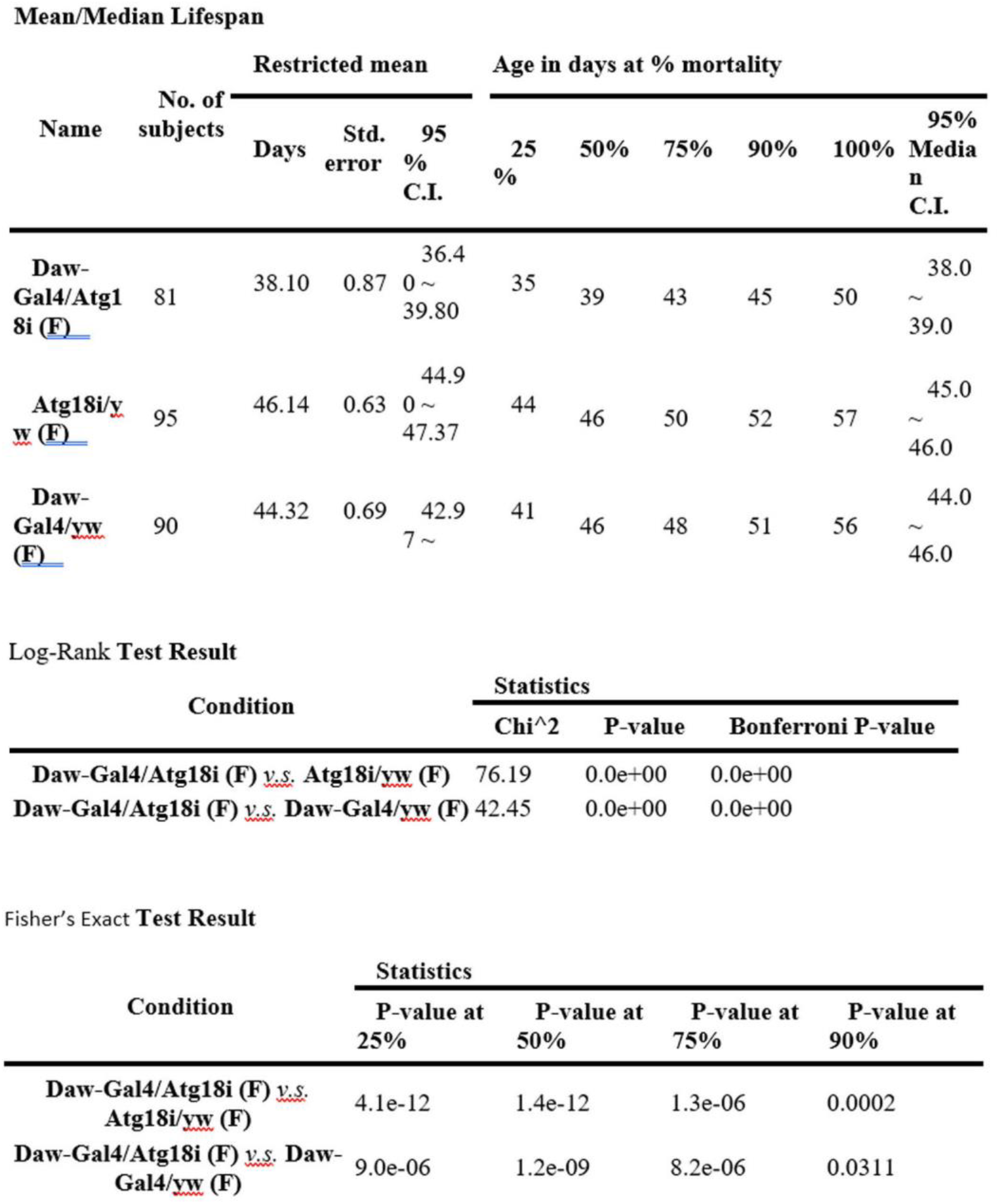
Statistical data on the lifespans of *Daw-Gal4* induced adult Atg18 knockdown female flies (*Daw Gal4/Atg18i (F)*) and their controls (*Atg18i/yw (F)* and *Daw-Gal4/yw (F)*) cultured at 29°C

**Supplemental table 12.**
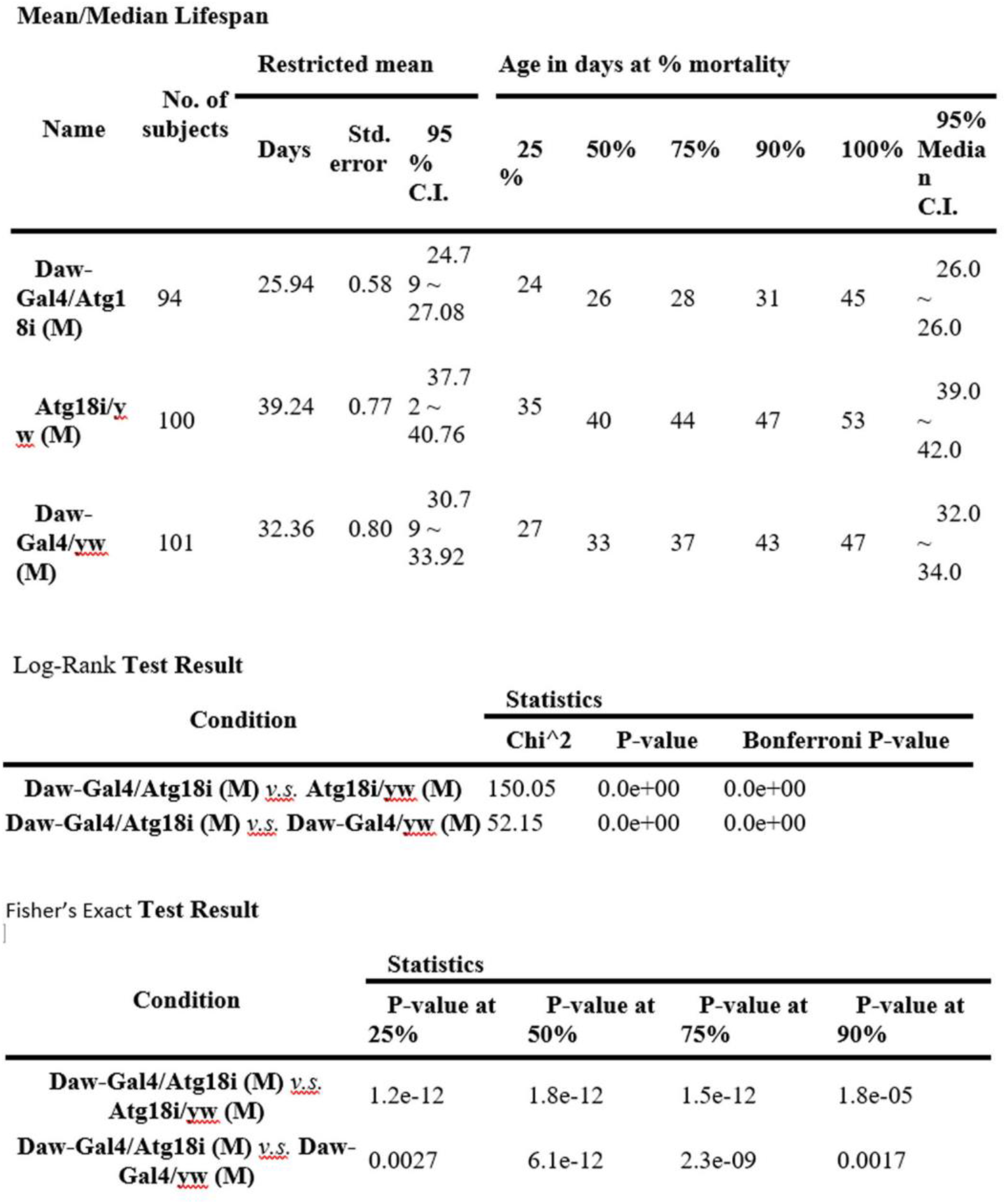
Statistical data on the lifespans of *Daw-Gal4* induced adult Atg18 knockdown male flies (*Daw Gal4/Atg18i (M)*) and their controls (*Atg18i/yw (M)* and *Daw-Gal4/yw (M)*) cultured at 29°C

